# Early cytokine-driven adaptation of survival pathways in lymphoid cells during targeted therapies

**DOI:** 10.1101/2024.03.04.583422

**Authors:** Meng-Xiao Luo, Tania Tan, Marie Trussart, Annika Poch, Thi Minh Hanh Nguyen, Terence P. Speed, Damien G. Hicks, Esther Bandala-Sanchez, Hongke Peng, Stéphane Chappaz, Charlotte Slade, Daniel T Utzschneider, Andreas Strasser, Rachel Thijssen, Matthew E Ritchie, Constantine S Tam, Geoff Lindeman, David CS Huang, Thomas E Lew, Mary Ann Anderson, Andrew W Roberts, Charis E Teh, Daniel HD Gray

## Abstract

Venetoclax, a first-in-class BH3 mimetic drug targeting BCL-2, has improved outcomes for patients with chronic lymphocytic leukemia (CLL). Early measurements of the depth of the venetoclax treatment response, assessed by minimal residual disease, are strong predictors of long-term clinical outcomes. Yet, there are limited data concerning the early changes induced by venetoclax treatment that might inform strategies to improve responses. To address this gap, we conducted longitudinal mass cytometric profiling of blood cells from patients with CLL during the first two months of venetoclax monotherapy. At baseline, we resolved CLL heterogeneity at the single-cell level to define multiple subpopulations in all patients distinguished by proliferative, metabolic and cell survival proteins. Venetoclax induced significant reduction in all CLL subpopulations coincident with rapid upregulation of pro-survival BCL-2, BCL-XL and MCL-1 proteins in surviving cells, which had reduced sensitivity to the drug. Mouse models recapitulated the venetoclax-induced elevation of survival proteins in B cells and CLL-like cells that persisted *in vivo*, with genetic models demonstrating that extensive apoptosis and access to the B cell cytokine, BAFF, were essential. Accordingly, analysis of patients with CLL that were treated with a different targeted therapy, the anti-CD20 antibody obinutuzumab, also exhibited marked elevation of BAFF and increased pro-survival proteins in leukemic cells that persisted. Overall, these data highlight the rapid adaptation of CLL cells to targeted therapies via homeostatic factors and support co-targeting of cytokine signals to achieve deeper and more durable long-term responses.

**Key points:**

- Leukaemic cells rapidly adapt to targeted therapy by elevating pro-survival protein expression.
- Cell attrition and increased bioavailability of homeostatic cytokines drive this heightened survival, highlighting avenues for more potent combination therapies.

## Introduction

Chronic lymphocytic leukemia (CLL), the most prevalent adult leukemia in many developed countries, is characterized by the accumulation of long-lived CD5^+^CD19^+^ B cells in the blood, bone marrow and secondary lymphoid organs. Most CLL cells are quiescent^1^, with a small population of proliferating cells responding to growth factor signals detectable in lymph nodes and blood^2^. Defective apoptosis is a hallmark of CLL mediated by elevated expression of the pro-survival protein, BCL-2^3^. Venetoclax (ABT-199), a small molecule inhibitor of BCL-2, is a highly efficacious treatment for CLL either as monotherapy^4,5^ or in combination with other agents^6–9^. Venetoclax induces rapid apoptosis, with the majority of susceptible cells dying within 4–8 hours of exposure *in vitro* and *in vivo*^10^. Administration of high concentrations of venetoclax in patients with bulky disease can cause tumor lysis syndrome^4^, so a weekly dose ramp-up regimen is typically used.

Despite the high efficacy of venetoclax in CLL (and acute myeloid leukemia (AML)), incomplete responses and emergent therapeutic resistance remain important challenges. Studies of patients who relapse after long-term venetoclax therapy have revealed multiple, non-mutually exclusive drug resistance mechanisms: (i) mutations in the binding groove of BCL-2^11–13^, (ii) overexpression of other members of the BCL-2 family pro-survival proteins not targeted by venetoclax (e.g. BCL-XL and MCL-1)^11,14^, or (iii) loss of BAX^15,16^ and/or TP53 dysfunction^17^. Yet how leukemic cells initially evade venetoclax-induced apoptosis and the setting in which resistance emerges remain unclear.

Here, we sought to address these questions by longitudinal deep profiling of blood cells from patients during the ramp-up period. We found striking elevation of pro-survival BCL-2 protein levels in residual CLL cells post-treatment that was recapitulated in murine models. We found evidence that heightened BCL-2 in surviving CLL cells was due to both the preferential loss of cells with relatively lower BCL-2 expression and cytokine-driven upregulation of BCL-2. Abrogation of BAFF-R signaling blocked venetoclax-induced BCL-2 upregulation *in vivo* in normal B cells. Moreover, obinutuzumab, a targeted therapy that kills CLL cells in a BCL-2-independent manner, also resulted in BCL-2 upregulation in persisting cells, accompanied by marked elevation of serum BAFF and APRIL. These findings highlight a mechanism by which potent B cell depleting therapies alleviate cell competition for cytokine-mediated survival signals in CLL cells, suggesting new avenues to achieve deeper responses.

## Methods

### Patient samples

Patients treated with venetoclax monotherapy (VENICE study; NCT02980731^18^), venetoclax-ibrutinib (CAPTIVATE; NCT02910583^19^) or combination time-limited venetoclax-obinutuzumab^7^ were recruited from the Department of Clinical Haematology Royal Melbourne Hospital and Peter MacCallum Cancer Centre (Victoria, Australia) (**Supplementary Table 1**); healthy donors were from the Victorian Blood Donor Registry. All donors provided written informed consent and the research was approved by Human Research Ethics Committees/Institutional Review Boards. PBMCs were isolated by density gradient centrifugation and resuspended in IMDM medium with 10% heat-inactivated foetal calf serum. Patients with breast cancer (#ACTRN12615000702516^20^, **Supplementary Table 1**) were analyzed by CITE-seq as previously described^21^.

### Mass cytometry

Cells were prepared as previously described^21,22^ and frozen at -80°C. Thawed cisplatin-labelled cells underwent 20-plex palladium barcoding and were batched (**Fig S1A**) with common anchor samples. After methanol permeabilization and staining with antibody conjugates (**Supplementary Table 2**) and 125nM ^191^Ir/^193^Ir DNA intercalator (Fluidigm, CA, USA), cells were washed, filtered and resuspended with EQ normalization beads and acquired on a Helios CyTOF (Fluidigm, CA, USA). Data were processed using the CATALYST package^23^. Replicated anchor samples were used to identify and correct for batch effects using CytofRUV^24^. FlowSOM with 22 lineage proteins was used to identify clusters and all clusters were used to define pseudo-replicates with k=3. Data were visualized using uniform manifold approximation and projection (UMAP)^25^.

### Linear Discriminant Analysis (LDA)

Linear discriminant analysis (LDA) was performed on IgM^+^CLL clusters for each patient with increasing doses of venetoclax. Results were displayed and ranked according to the magnitude of the projection of each marker direction onto the plane of the first two LDA components. The green shading represents the distribution of ranking curves when randomly picking a 2D plane in *n*-dimensional space (where *n* is the number of parameters), projecting each of the *n* markers onto it, then ranking the markers based on their projected length on that plane. This process was repeated 500 times and the standard deviation of the sampled projection lengths at each rank position is determined.

### *In vitro* cell death assays

Cells were treated with graded concentrations of venetoclax or dimethyl sulfoxide (DMSO). Cell viability was quantified by flow cytometry with propidium iodide (5μg/mL; Sigma).

### Cell counts

Cell number calculations by flow cytometry were performed using Green Live/Dead Cell Dye (Invitrogen) for viability. Concentration of cells in blood = % total live WBC (and daughter gates) by flow cytometry **×** WBC concentration in blood by Advia = *y* (cells/L)

### Mice

C57BL/6, *Bak*^-/-^*Bax*^ΔCd23^, *vav-huBcl2*, *Bak*^-/-^*Bax*^ΔCd23^*Tnfrsf13c*^-/-^ and *Eμ-TCL-1* mice were backcrossed onto the C57BL/6 background from foundation strains (**Supplementary Table 5**). All mice were housed at WEHI or University of Melbourne in SPF conditions and experiments performed under relevant Animal Ethics Committee requirements. Wild-type mice received 5×10^5^ splenocytes from leukemic *Eµ-TCL1* donors (%CD19^+^CD5^+^ cells>95%) by intravenous injection. Recipients were treated with venetoclax or vehicle after reaching 80% CD19^+^CD5^+^ of blood lymphocytes. Venetoclax (100mg/kg body weight) or vehicle was administrered by daily oral gavage for 1 week.

### Hematopoietic reconstitution

CD45.1/CD45.2 C57BL/6 F1 mice (6-8 week-old) were lethally irradiated with 2×5.5 Gy 3h apart and reconstituted by intravenous injection of 2×10^6^ bone marrow (BM) cells. Irradiated recipients received anti-Thy1 (clone T24; 100 μg) intraperitoneally 24h later then reconstituted for >6 weeks.

### Flow cytometry

Single-cell suspensions were stained with antibody conjugates to detect cell surface and intracellular proteins (**Supplementary Table 6**). Samples were acquired on LSRII or Fortessa flow cytometer (BD Bioscience) and analyzed using Flowjo software (Treestar).

### Cytokine measurements

Serum cytokines were measured using the Luminex xMAP technology on the Bio-Plex 200 platform (Bio-plex Manager 5.0) with the Bio-Plex Pro™ Human Cytokine Screening Test Kit (48-Plex, Bio-Rad) and the Bio-Plex Pro™ Human Inflammation Panel-1 Kit (37-Plex, Bio-Rad).

### Quantification and statistical analysis

Data were analyzed using GraphPad Prism software. Student’s two-tailed *t* tests (paired or unpaired) were used for comparisons and differences were considered significant when p<0.05.

## Results

### Mass cytometry resolves heterogeneity among CLL cells

To assess the impact of venetoclax on CLL heterogeneity and survival, we assayed peripheral blood (PB) from 18 relapse/refractory CLL patients receiving venetoclax monotherapy^18^, healthy donor and two treatment naïve CLL patients receiving venetoclax-ibrutinib after ibrutinib run-in^7^ (**Supplementary Table 1**). PB samples were drawn before each weekly venetoclax dose-escalation until reaching 400 mg/day (**Fig 1A**). As expected, venetoclax induced substantial dose-related reductions in PB lymphocytes in all patients (**Fig 1B**). Samples were palladium-barcoded and pooled into batches that included shared “anchor” samples then stained with a panel of 43 conjugates to discriminate CLL/normal immune sub-populations, BCL-2 family proteins, cell cycle regulators, key signaling pathways, including NF-κB, ERK/p38, mTOR, DNA damage, CREB and cancer-related proteins (TP53 and c-MYC)^22^ (**Fig S1A**, **Supplementary Table 3)** for CyTOF.

**Figure 1.**
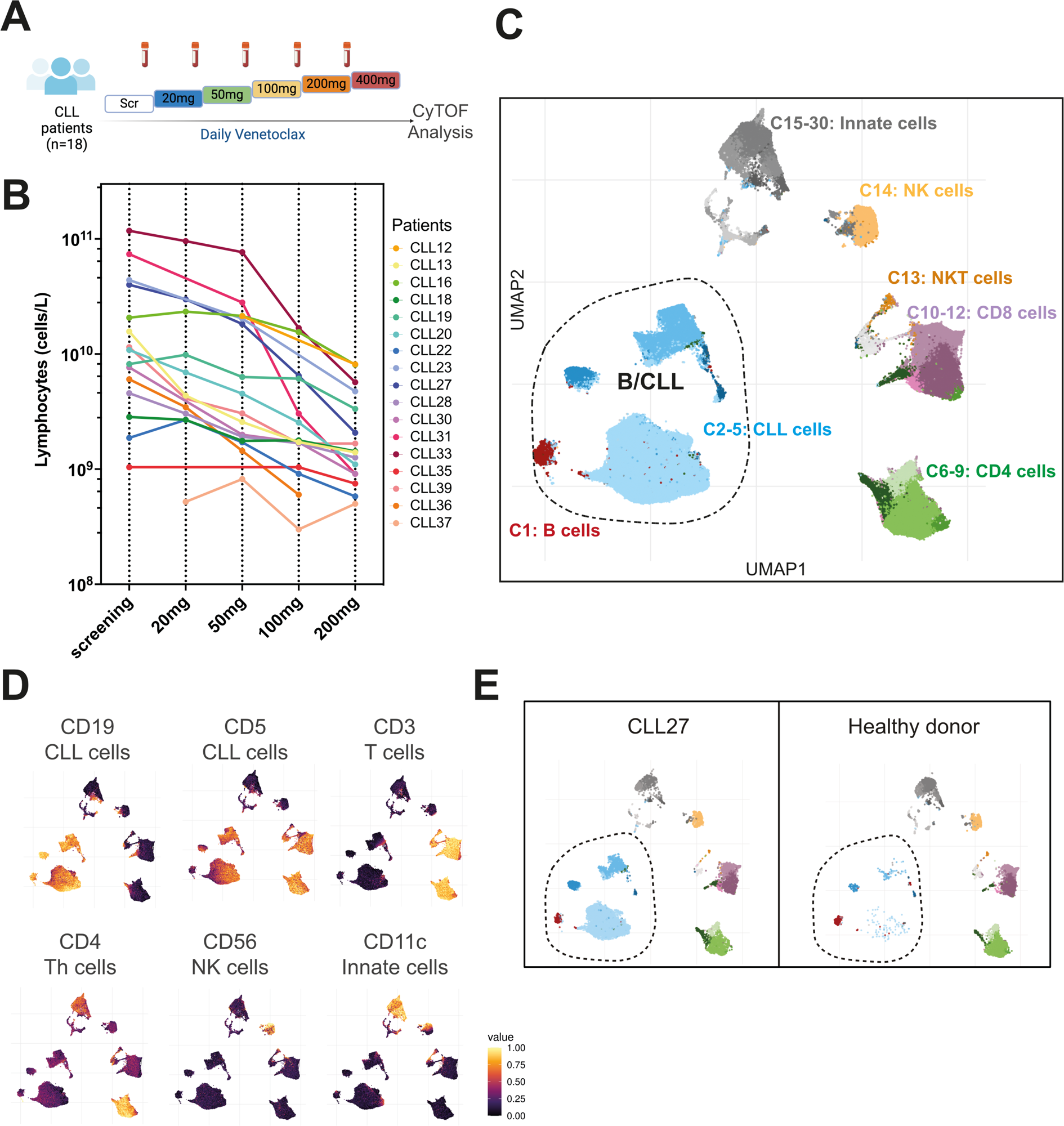
Mass cytometric analysis resolves CLL heterogeneity. **(A)** Schematic representation of the experimental strategy, with PB samples from 18 CLL patients collected at screening and during weekly venetoclax dose escalation. **(B)** Lymphocyte counts (cells/L) during venetoclax dose escalation in each patient. (**C**) UMAP projection of PB cells (subsampling of 2000 cells per each sample) from CLL patients (n=18 VEN patients plus n=2 VEN+IBR patients, 6 timepoints) and healthy donors coloured by cluster: B cells (C1), CLL cells (C2-5), T cells (C6-C12), NK cells (C14) and myeloid cells (C15:C30). **(D)** UMAP plots colored by median cell surface protein expression of indicated markers used to identify major immune cell populations. **(E)** UMAP plots of a patient with CLL (CLL27) and healthy donor with B/CLL clusters circled.

FlowSOM was used to identify 30 clusters and the data visualized with 2D UMAP (**Fig 1C**). These clusters were annotated based on the expression of lineage markers distinguishing healthy B (C1: CD19^high^CD5^low^), CLL (C2-5: CD19^high^CD5^high^), T (C6-12: CD3^high^), NKT (C13), NK (C14: CD56^high^) and myeloid (C15-30: CD11c^high^) cell populations (**Fig 1C-D**). Cluster distinctions within lineages were driven by expression of BCL-2 family proteins, cell cycle regulators, core signaling pathway constituents, including NF-kB, ERK/p38, mTOR, JAK/STAT and the transcription factors, pCREB, TP53 and c-MYC (**Fig S1B**). Overall, the initial analysis identified four putative CLL clusters (C2-5: CD19^high^CD5^high^) that were prominent in PB from CLL patients but not in a healthy donor (**Fig 1E**).

Next, we performed finer mapping of normal B/CLL clusters into 10 sub-clusters using FlowSOM (**Fig 2A**). Overlaying the expression profile of various markers (**Fig S1C)** on UMAPs distinguished normal CD5^low^CD20^high^pS6^high^ healthy B cells (C2.1-2.3)^26^ (**Fig 2B**) from CLL cells. CLL clusters could be broadly divided into IgM^low^ (C2.4-2.5) and IgM^high^ cells (C2.6-2.10) (**Fig 2C**). In addition, we detected minor CLL sub-populations, including pRB^high^ TACI^high^ BAX^high^ (C2.9) and pH2AX^high^ (C2.10) subsets (**Fig. 2A, 2D**).

**Figure 2.**
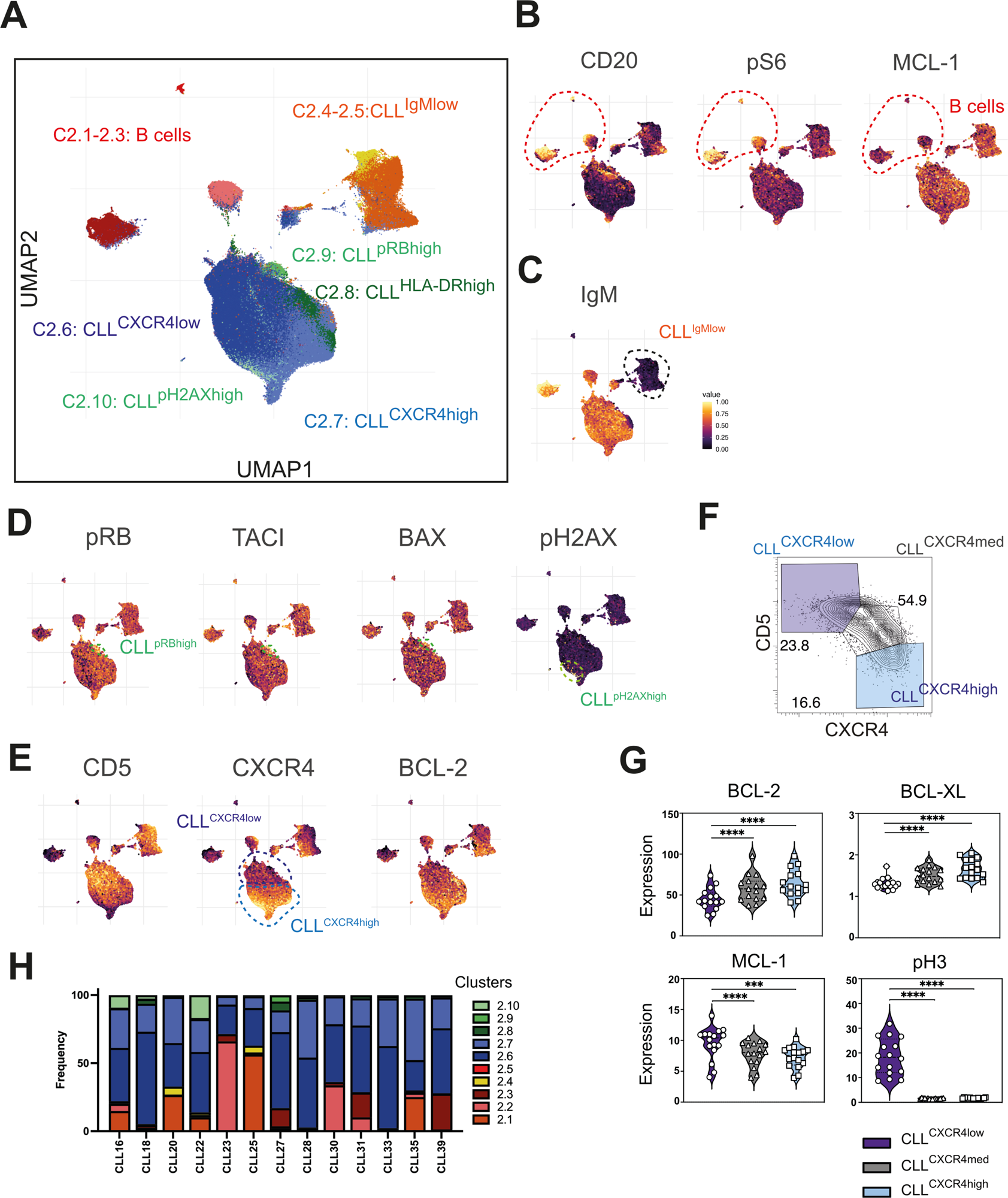
Intra- and inter-patient heterogeneity of CLL cell sub-populations. **(A)** UMAP of re-clustered healthy B/CLL cells (from **Fig 1C**) from patients and healthy controls. **(B)** UMAP plots colored by median protein expression of CD20, pS6 and MCL-1 distinguishing healthy B cells (CD20^high^ pS6^high^ MCL-1^low^) and CLL cells (CD20^low^pS6^low^MCL-1^high^). **(C)** UMAP plots colored by median protein expression of IgM, distinguishing IgM^high^ and IgM^low^ B cells. **(D)** UMAP plots colored by median amounts of pRB, TACI, BAX and pH2AX distinguishing two minor sub-populations of CLL cells: pRB^high^TACI^high^BAX^high^ (Cluster 2.9) and pH2AX^high^ (Cluster 2.10). **(E)** UMAP plots colored by median protein expression of CD5, CXCR4 and BCL-2, distinguishing two main sub-populations of CLL cells: CD5^high^CXCR4^low^BCL-2^high^ (Cluster 2.6) and CD5^low^CXCR4^high^BCL-2^v. high^ (Cluster 2.7). **(F)** Representative Flowjo plot of CXCR4 *vs* CD5 expression showing CD5^high^CXCR4^low^ and CD5^low^CXCR4^high^ CLL cell populations from patient CLL20. **(G)** Violin plots showing mean intensities of BCL-2, BCL-XL, MCL-1 and pH3 of old and new CLL cells. **(H)** Frequency of each cluster in PB of patients prior to venetoclax treatment. Student’s paired *t* test was used, *, **, *** and **** denotes P<0.05, P<0.01, P<0.005, P<0.001, respectively, n.s. denotes not significant.

Interestingly, the major CLL clusters had graded expression of BCL-2, CD5 and CXCR4 (**Fig 2E)**. Traditional 2D gating identified CXCR4^low^CD5^high^ and CXCR4^high^CD5^low^ CLL populations (**Fig 2F**), previously identified as PB CLL cells that have recently migrated from or are *en route* to lymphoid tissues, respectively^27^. Consistent with prior observations^28^, CXCR4^low^CD5^high^ CLL cells had relatively lower levels of BCL-2 and BCL-XL than CXCR4^high^CD5^low^ CLL cells, but higher amounts of MCL-1 and mitosis-associated phosphorylated histone 3 (pH3) (**Fig 2G**). With a few exceptions, all CLL clusters were found in all patients, albeit at varying frequencies (**Fig 2H**), suggesting relatively modest inter-patient variability. Taken together, the CyTOF profiling of CLL cells revealed heterogeneity distinguished by cell surface phenotype, signaling states (e.g. pRB, pH2AX) and expression of BCL-2 pro-survival proteins (e.g. BCL-2^high^CXCR4^high^CD5^low^ *vs* BCL-2^low^CXCR4^low^CD5^high^ CLL cells).

### Venetoclax induces a dose-related increase in BCL-2 in surviving CLL cells

Having defined the baseline characteristics and heterogeneity of CLL cell populations among patients, we sought to resolve the impact of venetoclax monotherapy (*n*=18). Venetoclax reduced CLL burden in all patients with no dose-dependent changes in CLL cluster proportions observed (**Fig 3A, Figure S2A**). Gating on CXCR4^low^CD5^high^ and CXCR4^high^CD5^low^ populations identified some instances of CXCR4^high^ CLL cell enrichment (with comparatively higher BCL-2), but this trend was not statistically significant (**Fig S2B, C**). To identify dose-related changes in CLL cells, independent of sub-populations, we performed linear discriminant analysis (LDA). LDA ranked proteins according to the most significant dose-dependent changes in CLL cells in individual patients (e.g. patient CLL19, **Fig 3B, Fig S3A**), compared to the distribution of randomly oriented planes (green shaded area). A summary of LDA from all patients showed that the top ranked dose-dependent change in CLL cells over time was BCL-2 (**Fig 3C**). Accordingly, we observed a significant increase in mean level of BCL-2 in CLL cells at post-50 mg and post-100 mg doses of venetoclax treatment compared to baseline (**Fig S3B**). These CyTOF data show that short-term venetoclax treatment enriched for or drove higher BCL-2 in surviving cells (hereafter termed VEN^surv^).

**Figure 3.**
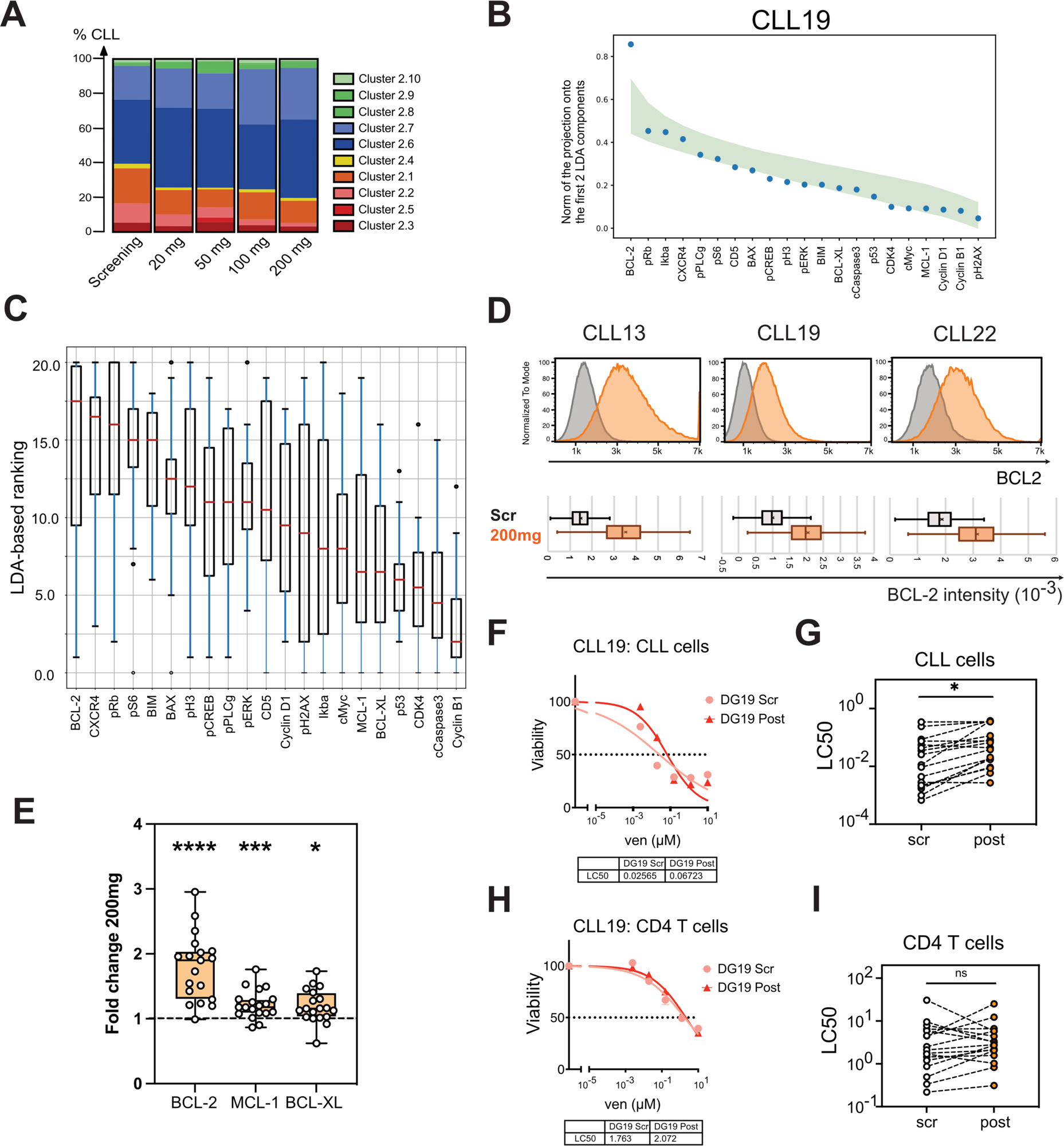
Venetoclax dose-dependent increase in BCL-2 protein detected in CLL cells. **(A)** Mean proportions of different CLL clusters from all patients in the cohort at indicated timepoints during treatment. **(B)** Graph of the results of linear discriminant analysis (LDA) of data from CLL clusters from patient CLL19 across increasing doses of venetoclax. The magnitude of the projection of each marker direction onto the plane of the first two LDA components is represented by blue dots. This has a maximum value of 1 if the marker direction lies in the planes of the first two LDA components. To reveal the markers driving the changes, the markers are ordered on the *x*-axis by the magnitude of this projection. The green shaded area represents the distribution of ranking curves of randomly oriented planes, see methods. (**C)** Summary of markers ranked by contributions to the first two LDA components in all patients for venetoclax dose escalation (seen in (B) for patient CLL19). The higher the ranking of a marker, the more it is affected by venetoclax dose changes, with 20 being the highest ranked marker and 0 the lowest. (**D)** Representative histograms showing mean BCL-2 protein expression of curated pairs of CLL patients at screening and after 200 mg VEN treatment in patient CLL13, CLL19 and CLL22 measured by flow cytometry. The distribution of BCL-2 levels in each sample is represented in the box and whisker plots in the lower panel. Box represents the 25^th^ 50^th^ and 75^th^ percentiles of the population, with x marking the mean and whiskers representing the minimum and maximum values (excluding outliers). **(E)** Fold change of BCL-2 MCL-1 and BCL-XL protein expression in samples at post-200mg normalized to screening in individual patients. P values were calculated using raw data and student’s two-tailed paired *t* test. (**F)** Representative results of venetoclax sensitivity *in vitro* assay in CLL cells from patient CLL19. **(G)** Summary of LC50 values of CLL cells at screening and post-200 mg venetoclax treatment. **(H)** Representative results of venetoclax sensitivity *in vitro* assay in non-transformed CD4^+^ T cells from patient CLL19. **(I)** Summary of LC50 values of CD4^+^ T cells at screening and post-200 mg venetoclax treatment. For box and whisker plots in D, outliers were omitted from the plot with an outlier factor of 1.5. Outliers represented less than 2.5% of total population in all samples, For G, I, Student’s two-tailed paired *t* test was used (patients with missing values were excluded from analysis). For E, G and I, *, **, *** and **** denotes P<0.05, P<0.01, P<0.005, P<0.001, respectively, n.s. denotes not significant.

To verify these findings, we used conventional flow cytometry with fluorochrome conjugates that offer a greater dynamic range than CyTOF. Analysis of paired samples at screening and post-200mg of venetoclax validated the significant increase in BCL-2 protein expression in VEN^surv^ cells (**Fig 3D-E**). Smaller, statistically significant increases in BCL-XL and/or MCL-1 were also apparent (**Fig 3E**). We compared the *in vitro* venetoclax sensitivity of cells at screening and post-200mg treatment and found reduced sensitivity in CLL cells (**Fig 3F-G**), but not CD4^+^ T cells (**Fig 3H-I)**. These data align with a previous study of a smaller patient group^29^ and demonstrate venetoclax selects for or induces cells with higher expression of pro-survival BCL-2 proteins shortly after therapy with resultant reduced sensitivity to venetoclax. To extend these findings into a setting of combination therapy, we analyzed two CLL patients that received first-line ibrutinib lead-in followed by ibrutinib plus venetoclax, comparing cells just before and after venetoclax treatment. The addition of venetoclax substantially reduced WBC counts (**Figure S3C**) and VEN^surv^ CLL cells in both patients had increased BCL-2 expression and reduced *in vitro* sensitivity to venetoclax (**Figure S3D-E**). These data suggest that the rapid adaptation processes acting on VEN^surv^ CLL cells are pertinent to multiple settings.

### Preferential loss of CLL cells with relatively lower BCL-2 protein partially accounts for VEN^surv^ cell phenotype

To explore how venetoclax causes an apparent, rapid increase in pro-survival proteins in VEN^surv^ cells, we hypothesized that, *in vivo*, dose escalation may preferentially eliminate those cells with relatively lower amounts of BCL-2 (**Fig 4A**). To test this notion, we first set a baseline threshold delineating the 20% of cells with the highest amount of BCL-2 (BCL-2^top20^) versus the remaining 80% (BCL-2^lower80^) cells in each patient at screening. We then applied this threshold to the paired post-200mg data (**Fig 4B**). Calculation of the *ex vivo* concentration of the two populations revealed a mean 30-fold decrease in BCL-2^lower80^ cells while BCL-2^top20^ cells decreased by only 3.2-fold (**Fig 4C**). These data support the notion that during the venetoclax ramp-up, CLL cells with lower amounts of BCL-2 are preferentially killed. However, we also observed that VEN^surv^ cells often manifest levels of BCL-2 that exceeded the range measured prior to venetoclax administration (**Fig 3D, 4B**), suggesting another (non-mutually exclusive) scenario where cell extrinsic factors drive upregulation of BCL-2.

**Figure 4.**
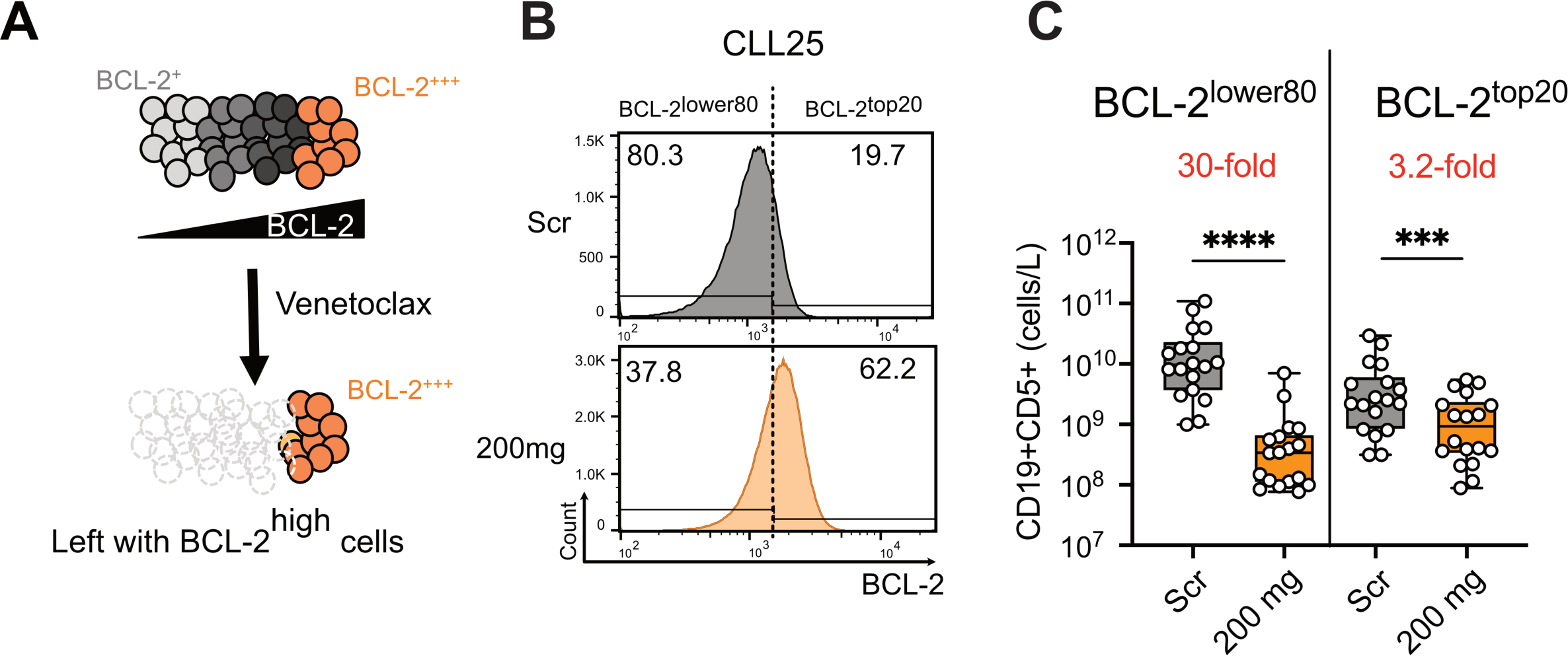
Preferential killing of CLL cells with relatively low BCL-2 levels partially accounts for the increase of BCL-2 in VEN^surv^ cells. (**A**) Schematic representation of the hypothesis: CLL cells express a range of BCL-2 amounts, from BCL-2^+^ to BCL-2^+++^. CLL cells with relatively low level of BCL-2 (BCL-2^+^) are sensitive to venetoclax treatment, enriching for CLL cells with higher BCL-2 levels (BCL-2^+++^) after venetoclax monotherapy. **(B)** Distribution of BCL-2 in CLL cells before (Scr) and one week after 200 mg dose of venetoclax with a dashed line indicating the threshold distinguishing the top 20% of BCL-2 expressors (BCL-2^top20^) and 80% lower BCL-2 expressors (BCL-2^lower80^) at screening. This threshold was then applied to each paired post-200 mg profile. **(C)** Estimated concentration of BCL-2^top20^ and BCL-2^lower80^ in CLL cells in the blood at screening and post-200 mg venetoclax treatment based on the BCL-2^top20^/BCL-2^lower80^ threshold assigned in the screening sample. For bar graphs, mean±SEM is shown, ratio paired *t* test was used, *, **, *** and **** denotes P<0.05, P<0.01, P<0.005, P<0.001, respectively.

### *In vivo* elevation of BCL-2 in VEN^surv^ cells in mice

To further dissect the mechanisms driving increased pro-survival protein expression in VEN^surv^ cells, we assayed normal and leukemic B cells in mice shortly after venetoclax treatment. Consistent with previous studies^30^, venetoclax swiftly reduced total B cell numbers by >2-fold in wildtype (wt) mice (**Fig 5A,B**), predominantly in peripheral B cells. As observed among CLL patients (**Fig 3E**), VEN^surv^ B cells had elevated levels of BCL-2 and MCL-1; BCL-XL levels remained unchanged (**Fig 5C-D**). In *Bak*^-/-^*Bax^ΔCd23^* mice, B cells lack the downstream effectors of apoptosis, BAX and BAK, therefore cannot undergo apoptosis. Venetoclax did not affect B cell numbers or BCL-2 expression in these mice (**Fig 5B-D**), demonstrating that apoptotic death was essential for elevation of BCL-2 in VEN^surv^ B cells. Intriguingly, MCL-1 levels still increased in VEN^surv^ B cells from *Bak*^-/-^*Bax^ΔCd23^* mice (**Fig 5B-D**), suggesting a non-apoptotic or cell extrinsic mechanism influences MCL-1 protein expression in this scenario.

**Figure 5.**
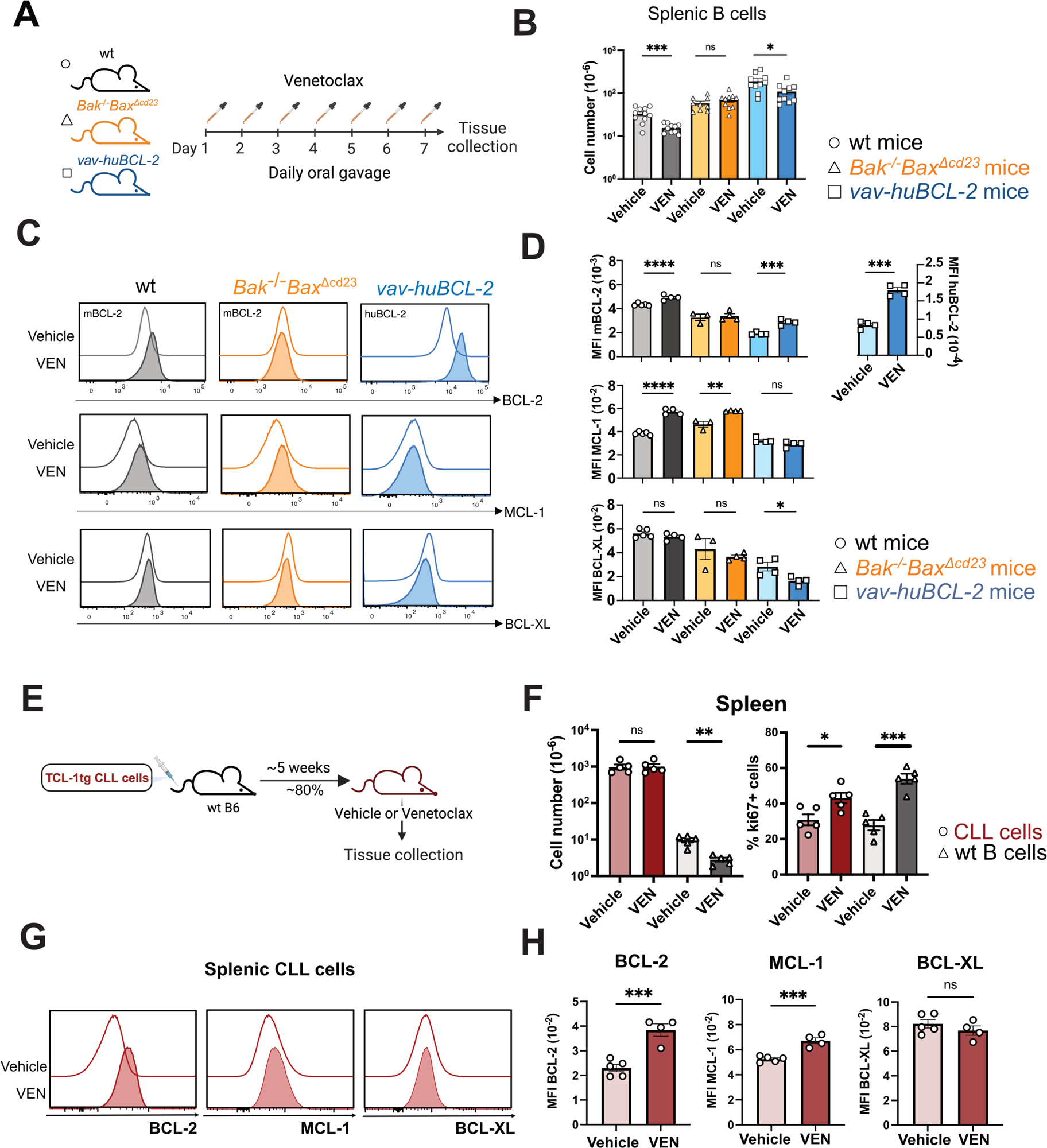
*In vivo* modelling demonstrates the elevation of BCL-2 in VEN^surv^ cells in mice. **(A)** Schematic representation of experimental strategy. Wildtype (wt), *Bak*^-/-^*Bax*^Δcd23^ and *vav-huBcl-2* mice were treated daily with 100 mg/kg body weight venetoclax or vehicle (a mixture of 60% Phosal 50 PG, 30% PEG 400 and 10% EtOH) for one week. **(B)** Absolute numbers of splenic B cells before and after venetoclax treatment in wt, *Bak*^-/-^*Bax*^Δcd23^ and *vav-huBcl-2* mice. **(C)** Histograms of BCL-2, MCL-1 and BCL-XL protein in CD19^+^CD21^+^IgM^+^ B cells from wt, *Bak*^-/-^*Bax*^Δcd23^ or *vav-huBcl-2* mice measured by flow cytometry after vehicle or venetoclax treatment. **(D)** Geometric mean (±SEM) of mBCL-2, huBCL-2, MCL-1 and BCL-XL protein levels in vehicle and venetoclax treated mice of the indicated genotypes. **(E)** Schematic representation of *E*μ*-TCL-1* transgenic mice modelling of CLL cell response to venetoclax. Cohorts of C57BL/6 mice received 5 x 10^5^ *E*μ*-TCL-1* transgenic CLL cells from the same donor, and once reaching 80% leukemic burden in blood, were treated with vehicle or venetoclax daily for 7 days. **(F)** Total splenic cell numbers and the proportions of Ki67^+^ *E*μ*-TCL-1* transgenic CLL cells and wt B cells before and after treatment. **(G)** Histograms of BCL-2, MCL-1 and BCL-XL protein in CLL cells recovered from the spleen of mice treated with vehicle or venetoclax. **(H)** Quantification of BCL-2, MCL-1 and BCL-XL protein expression in CLL cells recovered from the spleen of mice treated with vehicle or venetoclax. Data from (B-D) are representative of three independent experiments with n=2-5 mice per group. Data from (F-H) are representative of three independent experiments with n=5-6 mice per group. For all bar graphs, mean±SEM are shown and each symbol represents an individual mouse; Student’s two-tailed *t* test was used, *, **, *** and **** denotes P<0.05, P<0.01, P<0.005, P<0.001, respectively, n.s. denotes not significant.

We also assessed the impact of venetoclax on *vav-huBCL-2* transgenic mice that overexpress human BCL-2 in all hematopoietic cells^31^. B cells from *vav-huBCL-2* mice remained sensitive to venetoclax-induced cell death *in vivo* (**Fig 5B**) and VEN^surv^ cells exhibited substantially increased amounts of both human (i.e. transgene encoded) and mouse BCL-2 (i.e. endogenous) (**Fig 5C-D**). Together, these data indicate that the increase in cellular BCL-2 protein following short-term venetoclax treatment *in vivo* is: (1) conserved between mouse and human BCL-2, (2) dependent on cells undergoing apoptosis, (3) independent of the BCL-2 antibodies used for detection, and (4) observed in B cells with varying baseline amounts of BCL-2.

Next, we tested whether the elevated BCL-2 observed in VEN^surv^ B cells also occurs in a mouse model of CLL. We adoptively transferred CLL cells from *Eμ-TCL-1* transgenic mice into wt mice and treated recipients with either vehicle or venetoclax (**Fig 5E**). Venetoclax caused a significant decline in splenic healthy B cells, but not in leukemic CD19^+^CD5^+^ cells (**Fig 5F**). The apparently lower sensitivity of CD19^+^CD5^+^ *Eμ-TCL-1* transgenic B cells, consistent with previous studies^32,33^, is likely due to the high proliferation of *Eμ-TCL-1* CLL-like cells compared to human CLL cells. Indeed, the proportions of KI-67^+^ proliferating *Eμ-TCL-1* transgenic B cells increased with venetoclax treatment (**Fig 5F**), presumably maintaining their numbers despite substantial apoptosis. Nevertheless, we observed increased expression levels of BCL-2 and MCL-1 in VEN^surv^ leukemic cells (**Fig 5G-H**). Collectively, these data reveal that venetoclax-induced apoptosis, *in vivo*, rapidly drives elevated BCL-2 and MCL-1 protein levels in VEN^surv^ cells, recapitulating the responses observed in CLL cells from patients.

### Competition for BAFF controls the homeostatic response to venetoclax in murine B cells

We next sought to determine whether cell extrinsic factors drive the increase in BCL-2 proteins in VEN^surv^ cells. To establish an *in vivo* model where the impact of venetoclax on the microenvironment could be read-out in apoptosis-resistant B cells, we generated hematopoietic chimeras with a mixture of congenically-labelled wild-type (CD45.1^+^) cells and *Bak*^-/-^*Bax^ΔCd23^* (CD45.2^+^) cells into CD45.1/2 irradiated recipients (**Fig 6A**). These chimeras were treated with venetoclax which ablated wt B cells, accompanied by elevated BCL-2, MCL-1 and BCL-XL in surviving wt cells (**Fig 6B-D**). The numbers of B cells derived from *Bak^-/-^Bax*^ΔCd23^ BM was unchanged by venetoclax treatment; however, their levels of BCL-2 increased (**Fig 6B-D**). This finding contrasts the observations in intact *Bak^-/-^Bax*^ΔCd23^ mice (**Fig 5C-D**), indicating that BCL-2 upregulation is driven by cell extrinsic factors and dependent on the removal of wt cells in the chimeras. MCL-1 protein increased modestly in both wt and *Bak^-/-^Bax*^ΔCd23^ B cells from the venetoclax-treated chimeras (**Fig 6C-D**), similar to findings in intact *Bak^-/-^Bax*^ΔCd23^ mice (**Fig 5C-D**), indicating that the elevation of MCL-1 was independent of B cell apoptosis. Collectively, these data demonstrate that cell extrinsic cues rapidly upregulate pro-survival proteins in VEN^surv^ B cells.

**Figure 6.**
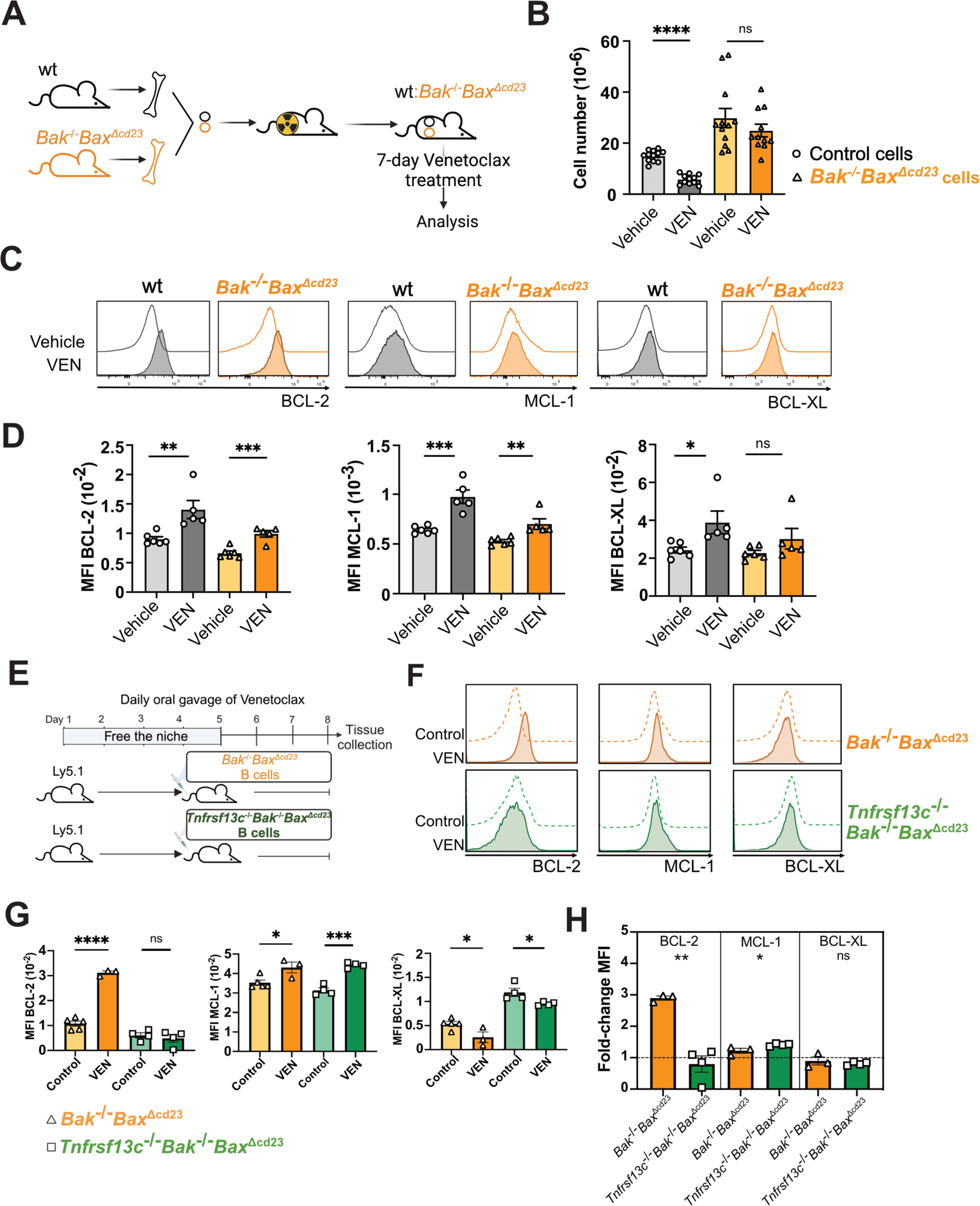
BAFF controls the homeostatic survival response to venetoclax treatment in B cells. **(A)** Schematic representation of hematopoietic chimeras reconstituted with a mixture of CD45.2^+^ *Bak*^-/-^*Bax*^Δcd23^ and CD45.1^+^ wt bone marrow progenitors, followed by venetoclax treatment. **(B)** Total numbers of splenic B cells of *Bak*^-/-^*Bax*^Δcd23^ or wt origin recovered from vehicle and venetoclax treated chimeras. **(C)** Histograms of BCL-2, MCL-1 and BCL-XL staining gated on splenic B cells of wt or *Bak*^-/-^*Bax*^Δcd23^ origin recovered from chimeric recipient mice treated with vehicle or venetoclax. **(D)** Geometric mean of BCL-2, MCL-1 and BCL-XL protein expression levels in splenic B cells of *Bak*^-/-^*Bax*^Δcd23^ or wt origin recovered from vehicle or venetoclax treated chimeric recipient mice. **(E)** Schematic representation of experimental design. Purified CD45.2^+^ C57BL/6 B cells from *Bak*^-/-^*Bax*^Δcd23^ or *Tnfrsf13c*^-/-^*Bak*^-/-^*Bax*^Δcd23^ mice were transferred into venetoclax treated CD45.1^+^ C57BL/6 wt recipient mice, which were then maintained on daily venetoclax treatment before analysis. **(F)** Histograms of BCL-2, MCL-1 and BCL-XL protein expression in donor CD45.2^+^ B cells from unmanipulated control mice with the same genotype or from venetoclax treated recipients. **(G)** Geometric mean of BCL-2, MCL-1 and BCL-XL protein expression in *Bak*^-/-^*Bax*^Δcd23^ and *Tnfrsf13c*^-/-^*Bak*^-/-^*Bax*^Δcd23^ B cells from control (untreated) recipient mice or from venetoclax treated recipient mice as indicated in (E), measured by flow cytometry. **(H)** Fold change of BCL-2, MCL-1 and BCL-XL protein levels in *Bak*^-/-^ *Bax*^Δcd23^ and *Tnfrsf13c*^-/-^*Bak*^-/-^*Bax*^Δcd23^ B cells in venetoclax treated mice. Data are expressed relative to the MFI in B cells recovered from unmanipulated control mice of the same genotype. Data from (B-H) are representative of three independent experiments with n=3-6 mice per group. For all bar graphs, mean±SEM are shown and each symbol represents an individual mouse; Student’s two-tailed *t* test was used, *, **, *** and **** denotes P<0.05, P<0.01, P<0.005, P<0.001, respectively, n.s. denotes not significant.

A prominent candidate cytokine is B cell activating factor (BAFF), which binds to the BAFF receptor (BAFF-R) on peripheral B cells to promote their maturation and survival^34^. To test this candidate, we used *Bak^-/-^Bax*^ΔCd23^ mice lacking the BAFFR (*Bak^-/-^Bax*^ΔCd23^*Tnfrsf13c*^-/-^)^35^, generating B cells that were both resistant to apoptosis and refractory to BAFF signaling. We adoptively transferred B cells purified from *Bak^-/-^Bax*^ΔCd23^ or *Bak^-/-^Bax*^ΔCd23^*Tnfrsf13c*^-/-^ mice into wt recipients that had been treated with venetoclax for four days to deplete their B cells (**Fig 6E**). After a further three-day treatment with venetoclax, we detected increased BCL-2 in *Bak^-/-^ Bax*^ΔCd23^, but not *Bak^-/-^Bax*^ΔCd23^*Tnfrsf13c*^-/-^ VEN^surv^ total B cells (**Fig 6F-H)** or IgM^+^CD21^+^ B cells (**Fig S4A-C**). By contrast, both populations showed modestly increased MCL-1 (**Fig 6F-H)**. These data demonstrate that venetoclax treatment rapidly induces BAFFR signals *in vivo* that upregulate BCL-2 in VEN^surv^ B cells, while a BAFF-independent pathway increases MCL-1 protein levels.

To assess whether a similar mechanism impacts human B cells, we investigated patients receiving venetoclax for metastatic breast cancer (i.e. without a hematological malignancy) (**Fig 7A**) (patient characteristics in **Supplementary Table 2**)^20^. Venetoclax treatment induced a striking reduction in circulating lymphocytes within 31 days (**Fig 7B**), consistent with the loss of circulating naïve B cells^20^. This decrease was accompanied by increased plasma concentrations of BAFF and APRIL (another cytokine mediating differentiation and survival of B cells) **(Fig 7C**) and elevated BCL-2 expression in B cells **(Fig 7D-E**). The impact of venetoclax treatment on the B cell transcriptome was analyzed using single cell CITE-seq data from 5 paired samples^21^. KEGG pathway analysis of DE genes (48 down, 67 up; false discovery rate <0.05, log fold change >0.5) showed 17 pathways enriched, including the BAFF-related NF-κB signaling, cytokine-cytokine receptor interactions, TNF signaling pathways and apoptosis (**Fig 7F**). These findings support the concept that increased BAFF signaling drives BCL-2 upregulation in VEN^surv^ B cells.

**Figure 7.**
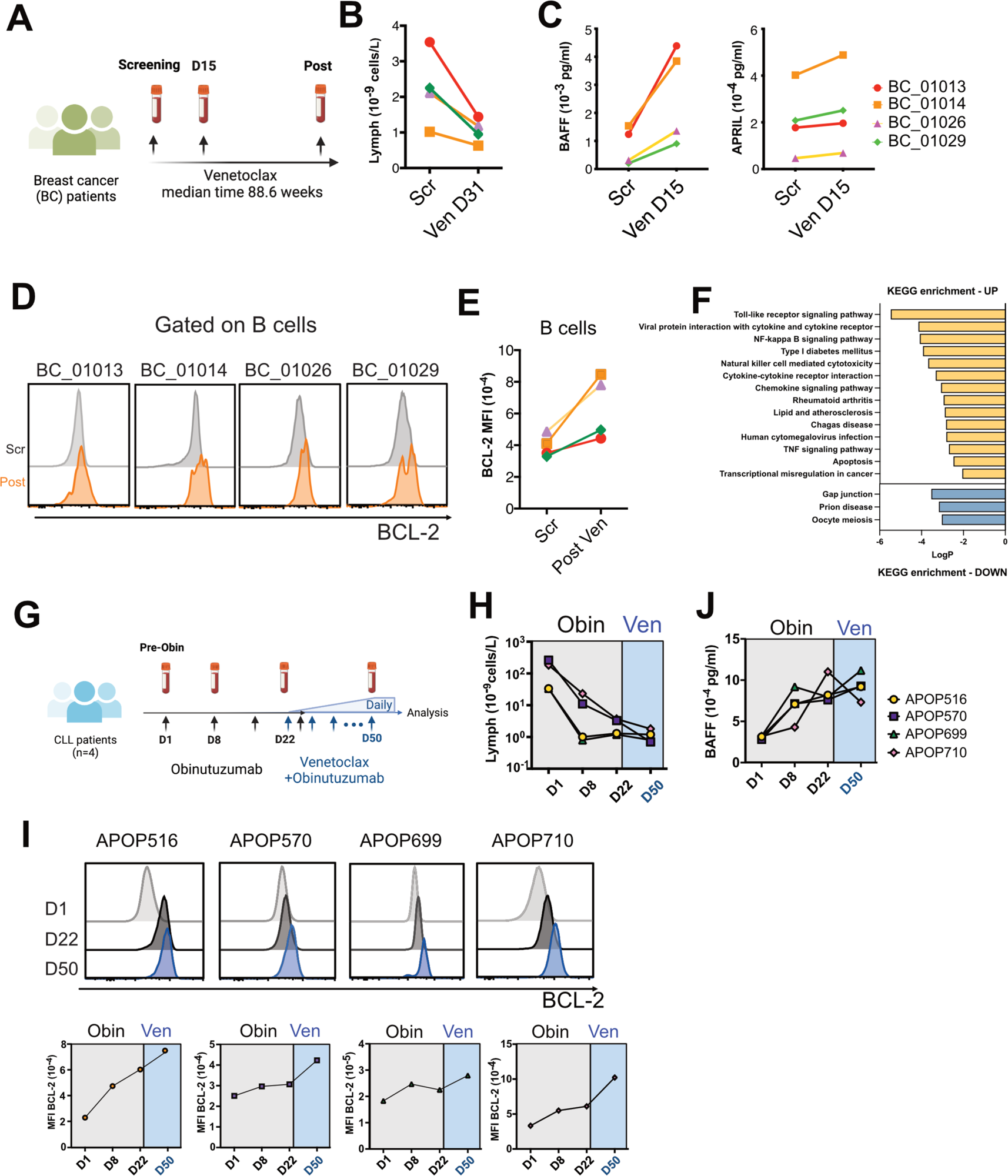
Increase in serum BAFF levels correlates with BCL-2 up-regulation in both healthy and malignant B cells in patients undergoing targeted therapies. **(A)** Schematic representation of study design. PB samples from 4 breast cancer patients were collected at screening and during venetoclax treatment. (all patients receiving 800 mg venetoclax) **(B)** Change of lymphocytes (cells/uL) during venetoclax treatment in each patient after 31 days of treatment. **(C)** Concentrations of BAFF and APRIL (pg/ml) detected in the serum samples of the breast cancer patients using Luminex assay at screening and 15 days after venetoclax treatment. **(D)** Histograms of BCL-2 protein expression by flow cytometry in B cells in patients before and after venetoclax treatment (BC_01013 after 201.7 weeks, BC_01014 after 61.1 weeks, BC_01026 after 67.4 weeks, BC_01029 after 112.1 weeks). **(E)** Geometric mean of BCL-2 levels of B cells in four patients. **(F)** Kyoto Encyclopedia of Genes and Genomes (KEGG) enrichments analysis of DE genes in single cell CITE-seq data from 5 paired samples under long-term venetoclax treatment. **(G)** Schematic representation of study design. PB samples from 3 CLL patients were collected at screening and during obinutuzumab and venetoclax dose escalation. Patients received intravenous infusion of obinutuzumab on days 1/2 (first 1000mg dose divided over two days 100/900 mg), 8 (1000 mg), 15 (1000 mg) and day 29 (1000 mg). Venetoclax was introduced at 20 mg daily on day 22 and increased weekly (20, 50, 100, 200, 400 mg). **(H)** Change of lymphocytes concentrations (cells/L) during obinutuzumab and venetoclax treatment in each patient. **(I)** Histograms (top panel) and MFI (lower panel) of BCL-2 protein expression in CLL cells in each patient at the indicated timepoints. **(J)** Concentration (pg/mL) of BAFF detected in the serum samples of the CLL patients using Luminex assay.

### Depletion of leukemia via other targeted therapies also elevates BCL-2 pro-survival proteins in surviving cells

The data thus far support the hypothesis that venetoclax-induced killing of leukemic cells reduces competition among VEN^surv^ cells for cytokines, such as BAFF, resulting in elevated BCL-2 protein expression and reduced sensitivity to venetoclax (**Figure S5A**). One prediction of this model is that any targeted therapy capable of sufficiently reducing leukemic burden would induce a similar effect in surviving CLL cells. We assayed CLL cells from four patients treated with anti-CD20 antibody (obinutuzumab) prior to venetoclax ramp-up during standard-of-care venetoclax-obinutuzumab therapy (**Fig 7G**, patient characteristics in **Supplementary Table 1**). Obinutuzumab monotherapy markedly reduced PB lymphocytes, with a further reduction in three patients upon subsequent venetoclax treatment (**Fig 7H**). BCL-2 expression increased in CLL cells surviving obinutuzumab monotherapy and increased further following venetoclax treatment (**Fig 7I**). We then measured serum cytokines in these patients on day 1, 8, 22 (obinotuzumab run-in) and 50 (post-venetoclax) (selected cytokines in **Figure S5B**). Within 8 days of obinutuzumab treatment, BAFF concentration increased ∼2-fold in all patients and remained high thereafter **(Fig 7J**). Together, these findings show that elevation of BCL-2 in cells surviving targeted therapies is not a unique feature of targeting BCL-2. Instead, it appears related to the reduction in leukemic cells and may reflect reduced cell competition and enhanced access to cytokines.

## Discussion

Venetoclax and other targeted therapies have improved treatment outcomes for patients with CLL and AML in certain settings; however, therapeutic resistance remains an important problem. It is clear that long-term venetoclax treatment in patients can induce a variety of changes in the interplay among BCL-2 family proteins that engender resistance^11–14,17^. Yet, a detailed understanding of how targeting BCL-2 over the short term impacts the apoptosis pathway within the context of CLL heterogeneity is lacking. This timeframe is important because: (1) patients with incomplete responses are more prone to relapse^4^ and (2) the prevailing environment supporting leukemic cell survival likely influences the manifestation of resistance. Our findings highlight the rapid elevation of BCL-2 in circulating CLL cells as a key feature in patients during venetoclax dose-escalation. We find evidence that, *in vivo*, BAFF-BAFF-R signaling contributes to BCL-2 upregulation, culminating in reduced sensitivity of CLL cells to treatment. Collectively, these data support the notion that alleviation of competition for BAFFR-mediated survival signals among CLL cells during treatment may limit therapeutic responses and support resistance.

Our deep profiling of circulating CLL cells at the single-cell level revealed that they could be further distinguished from normal B cells by lower expression of MCL-1 and pS6, consistent with their low metabolic activity and turnover. Among CLL cells, a key distinction was the expression of pro-survival proteins, demarcated by CXCR4 expression^36,37^. CXCR4^low^ cells, that had presumably recently emigrated from lymph nodes, were MCL-1^high^BCL-2^low^BCL-XL^low^ and proliferative (pH3^high^). By contrast, PB CXCR4^high^ CLL cells capable of migrating into lymph nodes were MCL-1^low^BCL-2^high^BCL-XL^high^ and quiescent (pH3^low^). These data reveal dynamic regulation of the expression of pro-survival BCL-2 family proteins in CLL cells associated with division.

All PB CLL clusters were diminished by venetoclax dose-escalation, concomitant with enrichment for cells with higher expression of BCL-2 and, to a lesser extent, BCL-XL and MCL-1. These data are consistent with a previous study of 5 CLL patients that found heightened expression of BCL-2 in surviving cells after 2-week venetoclax treatment^29^. We found that VEN^surv^ CLL cells exhibited reduced *in vitro* sensitivity to venetoclax and that the increased BCL-2 expression could only partially be explained by depletion of CLL cells with relatively lower BCL-2. *In vivo* mouse models provided clear evidence for a cell extrinsic mediator, transduced by BAFFR, that drove up pro-survival BCL-2 in VEN^surv^ B cells. Consistent with a mechanism whereby increased BAFF availability drives BCL-2 upregulation upon extensive CLL cell apoptosis, obinutuzumab treatment also led to elevated BCL-2 in a small patient cohort, coincident with markedly increased circulating BAFF. This phenomenon was also observed among two patients receiving venetoclax-ibrutinib after ibrutinib run-in, suggesting that cytokine-mediated BCL2 upregulation may also contribute to resistance to dual BCL2-BTK inhibition regimens, which are likely to be increasingly used in clinical practice^38,39^. These data support the notion that competition for BAFF constrains expression of pro-survival BCL-2.

We also detected modest increases in MCL-1 in surviving CLL cells, apparently driven by cell extrinsic signals that were independent of B cell apoptosis. One potential explanation is that MCL-1, which has a high rate of protein turnover, is stabilized in presence of venetoclax due to the displacement of pro-apoptotic BIM from BCL-2 to MCL-1^40,41^. Another possibility is the existence of an alternative pathway that upregulates MCL-1 in presence of venetoclax, perhaps via alteration of stromal cells^42^.

An implication of these data is that combining targeted therapies with agents that neutralize relevant cytokines could be beneficial by restraining BCL-2 expession. More specifically, our work builds the rationale to explore the combination of venetoclax with BAFF blockade. Indeed, co-targeting of BAFF with ibrutinib provided a survival benefit in a murine CLL model^43^ and the *in vitro* killing of primary patient CLL cells treated with venetoclax, idelalisib or ibrutinib^43,44^. A clinical trial combining venetoclax with a BAFF neutralizing antibody for treating CLL patients (NCT05069051) is currently in progress. Here we provide evidence for a potential *in vivo* mechanism-of-action, involving the prevention of BCL-2 protein upregulation and enhanced sensitivity to apoptosis. These findings raise the question of whether the elevated levels of homeostatic cytokines observed during targeted therapy purely reflect reduced consumption or involve microenvironmental responses to leukemic cell death.

More broadly, this cytokine/pro-survival protein axis may be relevant to other malignancies where venetoclax has been tested, such as AML and myeloma, where malignant cells utilize distinct homeostatic cytokines to support their survival. Together, our discoveries underscore the potential of mitigating the bioavailability of pro-survival cytokines when designing more potent treatment strategies in diverse types of hematologic cancers, which can induce deeper response and minimize the risk of future relapse.

## Acknowledgements

The authors thank the patients who enrolled in the venetoclax clinical trials and for assistance from Naomi Sprigg and Kelli Gray with the collection and curation of patient samples. In addition, the authors thank Andrew Mitchell and Tian Zheng at the Materials Characterisation and Fabrication Platform (MCFP) at the University of Melbourne for mass cytometry support.

This work was supported by grants and fellowships from the Australian National Health and Medical Research Council (NHMRC) fellowships (1089072) (C.E.T.), (1090236 and 1158024) (D.H.D.G.), (1078730 and 1175960) (G.J.L.), and (1043149 and 1156024) (D.C.S.H.); Investigator grants (1177718) (M.A.A.), (1194779) (D.T.U), (1174902) (A.W.R.); a Fulbright Australia-America Postdoctoral Fellowship (C.E.T.); Victorian Cancer Agency Fellowships (MCRF20026) (C.E.T.); Postdoctoral Fellowship from the German Cancer Aid (A.P.); Cancer Council of Victoria Grants-in-Aid (1146518 and 1102104) (D.H.D.G.) (D.T.U.); Perpetual Impact Philanthropy funding (C.E.T.); Australian NHRMC grants (2002618) (C.E.T.), (1016647, 1113577, 1016701, 1113133, 1079560, 2013478 and 2011139) (A.W.R. and D.C.S.H.), (1113133 and 1153049) (G.J.L.), (2013478) (D.C.S.H. and R.T.); The Medical Advances Without Animals Trust (MAWA), the Leukemia and Lymphoma Society, US, Specialized Center of Research grant (7015-18) (A.W.R., A.S., and D.C.S.H.); a Tour de Cure Mid-Career Research Grant (RSP-070-FY2023) (D.T.U.); National Breast Cancer Foundation Australia (IIRS-19-004); Breast Cancer Research Foundation (BCRF-20-182) (G.J.L.); Rae Foundation (V.L.B.); and Investigator Initiated Study support from AbbVie and Genentech (Roche) for the breast cancer study (#ACTRN12615000702516) (G.J.L).

This work was performed in part at the Materials Characterization and Fabrication Platform at the University of Melbourne and the Victorian Node of the Australian National Fabrication Facility with support from the Victorian Comprehensive Cancer Centre. This work was also supported by the Australian Cancer Research Foundation and made possible through Victorian State Government Operational Infrastructure Support and Australian Government NHMRC Independent Research Institutes Infrastructure Support Scheme.

