## Supplemental materials for "Early cytokine-driven adaptation of survival pathways in lymphoid cells during targeted therapies"

**List of Supplementary Materials:**

Supplementary Table 1. CLL Patient characteristics

Supplementary Table 2. Breast cancer Patient characteristics

Supplementary Table 3. Antibodies used for CyTOF analysis

Supplementary Table 4. Bioinformatic tools used for analysis

Supplementary Table 5. Drug treatment

Supplementary Table 6. Murine experimental model

Supplementary Table 7. Flow cytometry antibodies

Supplementary Figure 1.

Supplementary Figure 2.

Supplementary Figure 3.

Supplementary Figure 4.

Supplementary Figure 5.

**Supplementary Table 1. CLL patient characteristics.** Abbreviations: Chlorambucil – CLB; Rituximab – R; FR – fludarabine, rituximab; FCR – fludarabine, cyclophosphamide, rituximab; R-CHOP – rituximab, cyclophosphamide, doxorubicin, vincristine, prednisolone; CVP – cyclophosphamide, vincristine, prednisolone; BR – bendamustine, rituximab. IBR – ibrutinib; VEN – venetoclax; OBIN – Obinutuzumab; Best ever clinical response: PR – partial remission; CR – complete remission; MRD – Minimal residual disease by flow cytometry at $10^{-4}$, VEN+ IBR: Ibrutinib run-in followed by time-limited venetoclax-ibrutinib combination; VEN-O: Venetoclax-obinutuzumab standard of care.

| Patient ID | Age | Gender | Cytogenetic abnormality/mutation status (prior to commencing treatment) | | | | | Other therapies (prior to VEN treatment) | Agent | Best response on VEN | MRD | Progression free survival |
|---|---|---|---|---|---|---|---|---|---|---|---|---|
| | | | Chromosome 11q deletion | Chromosome 17p deletion | Complex karyotype | TP53 mutation | IGVH status | | | | | |
| CLL12 | 77 | F | Not detected | Not detected | Not detected | Not detected | Mutated | CLB; Navitoclax | VEN | PR | Not detected | Died in remission at 75 months |
| CLL13 | 74 | M | Not detected | Not detected | Not detected | Not detected | Unknown | CLB-R | VEN | PR | Detected | Disease progression at 72 months |
| CLL15 | 55 | M | Not detected | Not detected | Not detected | Not detected | Unmutated | None | VEN+IBR | CR | Detected | Alive and progression free at 79 months |
| CLL16 | 57 | M | Not detected | Detected | Detected | Detected | Unmutated | FCR; FCR re-treatment; ofatumumab | VEN | PR | Detected | Disease progression at 72 months |
| CLL18 | 64 | F | Not detected | Detected | Detected | Detected | Unknown | R; F-R; FCR | VEN | PR | Not detected | Disease progression at 17 months |
| CLL19 | 64 | M | Not detected | Not detected | Not detected | Not detected | Unmutated | R-CVP | VEN | CR | Not detected | Alive and progression free at 74 months |
| CLL20 | 59 | M | Not detected | Not detected | Not detected | Not detected | Unmutated | CVP; CHOP; FCR; CVP | VEN | PR | Detected | Alive and progression free at 75 months |
| CLL21 | 55 | M | Not detected | Not detected | Unknown | Unknown | Mutated | None | VEN+IBR | CR | Not detected | Alive and progression free at 75 months |
| CLL22 | 62 | M | Unknown | Unknown | Unknown | Unknown | Mutated | CLB; FCR | VEN | PR | Detected | Died in remission at 25 months |

| | | | | | | | | | | | |
|---|---|---|---|---|---|---|---|---|---|---|---|
| CLL23 | 70 | M | Not detected | Not detected | Unknown | Unknown | Unmutated | FCR | VEN | CR | Detected | Died in remission at 19 months |
| CLL25 | 52 | M | Detected | Not detected | Not detected | Not detected | Unmutated | FCR | VEN | PR | Not detected | Disease progression at 14 months |
| CLL27 | 62 | M | Detected | Detected | Detected | Detected | Unmutated | FCR; lenalidomide; BR | VEN | CR | Not detected | Disease progression at 35 months |
| CLL28 | 75 | F | Not detected | Not detected | Not detected | Not detected | Mutated | CLB; FCR; BR | VEN | CR | Not detected | Alive and progression free at 71 months |
| CLL30 | 58 | M | Not detected | Detected | Not detected | Not detected | Unmutated | Idelalisib- R | VEN | PR | Detected | Proceeded to allogeneic stem cell transplant at 21 months |
| CLL31 | 60 | M | Detected | Not detected | Detected | Detected | Unmutated | CLB; FCR; Lenalidomide ; Zanubrutinib | VEN | PR | Not detected | Disease progression at 11 months |
| CLL33 | 81 | M | Detected | Not detected | Not detected | Not detected | Unknown | R-CHOP | VEN | PR | Detected | Alive and progression free at 69 months |
| CLL35 | 78 | F | Unknown | Detected | Detected | Detected | Mutated | FCR; CLB-OBIN; IBR; R-methylpredni solone | VEN | Unknow n | Not detected | Died at 1 month |
| CLL36 | 67 | M | Detected | Not detected | Not detected | Not detected | Unmutated | R-CVP; rituximab; R-CVP; FCR; IBR | VEN | PR | Detected | Disease progression at 28 months |
| CLL37 | 46 | M | Not detected | Detected | Detected | Detected | Unmutated | FCR; idelalisib; IBR; BR | VEN | CR | Not detected | Proceeded to allogeneic stem cell transplant at 9 months |
| CLL39 | 71 | F | Unknown | Not detected | Unknown | Unknown | Mutated | R | VEN | CR | Not detected | Alive and progression free at 58 months |
| APOP516 | 55 | M | Detected | Not detected | Not detected | Not detected | Unmutated | None | VEN-O | CR | Not detected | Alive and progression free at 15 months |
| APOP570 | 79 | F | Not detected | Not detected | Not detected | Detected | Unmutated | None | VEN-O | Unkno wn | Unknown | Alive and progression free at 11 months |

| | | | | | | | | | | | | |
|---|---|---|---|---|---|---|---|---|---|---|---|---|
| APOP699 | 69 | M | Detected | Not detected | Not detected | Not detected | Unmutated | None | VEN-O | Unknown | Unknown | Alive and progression free at 6 months |
| APOP710 | 86 | M | Not detected | Not detected | Detected | Not detected | Unmutated | None | VEN-O | Unknown | Unknown | Alive and progression free at 4 months |

22

**Supplementary Table 2. Breast cancer patient characteristics**

| | ER % (strength 0 = none 1 = weak 2 = mod 3 = strong) | PR % (strength 0 = none 1 = weak 2 = mod 3 = strong) | BCL-2 % (strength 0 = none 1 = weak 2 = mod 3 = strong) | No previous therapy | Response |
|---|---|---|---|---|---|
| BC_01013 | 95 (3) | 95 (3) | 95 (3) | 0 | Partial response |
| BC_01014 | 80 (2) | 50 (2) | 90 (3) | 5 | Stable disease |
| BC_01026 | 90 (3) | 30 (2) | 90 (3) | 1 | Partial response |
| BC_01029 | 80 (3) | 70 (3) | 90 (3) | 0 | Partial response |

**Supplementary Table 3. Antibodies used for CyTOF analysis**

| Antibody | Clone | Metal Label | Source | Conc (ug) | Positive control | Negative control | RRID | Cat. |
|---|---|---|---|---|---|---|---|---|
| CD45 | HI30 | 89 Y | Biolegend | 2 | PBMC | NIH3T3 | AB_2562821 | 304045 |
| HLA-DR | L243 | 115 In | Biolegend | 2 | PBMC | PBMC | AB_314680 | 307602 |
| CD27 | M-T271 | 140 Ce | Biolegend | 4 | PBMC | PBMC | AB_2561786 | 356401 |
| CD235 | HIR2 | 141 Pr | Biolegend | 2 | NIH3T3 | PBMC | AB_2562825 | 306615 |
| CD19 | HIB19 | 142 Nd | Biolegend | 2 | PBMC | PBMC | AB_2562815 | 302247 |
| CD5 | UCHT2 | 143 Nd | Biolegend | 2 | PBMC | PBMC | AB_2563756 | 300627 |
| CD4 | RPA-T4 | 145 Nd | Biolegend | 2 | PBMC | PBMC | AB_2562809 | 300541 |
| IgM | MHM-88 | 146 Nd | Biolegend | 3 | PBMC | PBMC | AB_2563776 | 314527 |
| CD20 | H1 | 147 Sm | BD | 2 | PBMC | PBMC | AB_396030 | 555677 |
| CD16 | 3G8 | 148 Nd | Biolegend | 2 | PBMC | PBMC | AB_2562814 | 302051 |
| CD25 | 2A3 | 149 Sm | DVS | 1:100 | PBMC | PBMC | AB_2756416 | 3149010B |
| CD43 | 84-3C1 | 150 Nd | eBioscience | 2 | U937 | SUDHL4 | AB_763493 | 14-0439-82 |
| CD56 | HCD56 | 155 Gd | Biolegend | 2 | PBMC | PBMC | AB_2562830 | 318345 |
| CXCR4 (CD184) | 12G5 | 156 Gd | Biolegend | 2 | Jurkat | SUDHL4 | AB_2562824 | 306523 |
| CD10 | HI10a | 158 Gd | Biolegend | 2 | SUDHL4 | Jurkat | AB_2562828 | 312223 |
| CD11c | Bu15 | 159 Tb | Biolegend | 2 | MOLM13 | NB4 | AB_2562834 | 337221 |
| CD79b | CB3-1 | 162 Dy | BD | 2 | SUDHL4 | Jurkat | AB_396031 | 555678 |
| TACI (CD267) | 1A1 | 163 Dy | Biolegend | 3 | PBMC | PBMC | AB_528979 | 311902 |
| CD45RA | HI100 | 169 Tm | Biolegend | 2 | PBMC | PBMC | AB_2562822 | 304143 |
| CD3 | UCHT1 | 170 Er | Biolegend | 2 | PBMC | NIH3T3 | AB_2562808 | 300443 |
| CD38 | HIT2 | 172 Yb | Biolegend | 2 | MOLM-13 | MV4;11 | AB_2562819 | 303535 |
| CD61 | VI-PL2 | 209 Bi | DVS | 1:100 | PBMC | PBMC | AB_2864731 | 3209001B |
| Cyclin D1 | 92G2 | 139 La | CST | 4 | CLL irradiated | CLL | AB_2259616 | 2978S |

| | | | | | | | | |
|---|---|---|---|---|---|---|---|---|
| pPLCg2 [pY759] | K86-689.37 | 144 Nd | DVS | 1:100 | Ramos + anti-IgM | Ramos | NA | 3144015A |
| Cyclin B1 | GNS11 | 151 Eu | BD | 2 | PBMC | PBMC | AB_395290 | 554179 |
| pH3 | HTA28 | 152 Sm | Biolegend | 2 | PBMC | PBMC | AB_2562851 | 641007 |
| BCL-XL | E18 | 153 Eu | Abcam | 1 | HCT116 | REH | NA | ab199099 |
| BAX | 1B4 | 154 Sm | WEHI (Huang) | 2 | KMS-12-PE EV | KMS-12-PE sgBax | NA | NA |
| BCL-2 | 100 | 157 Gd | WEHI (Strasser) | 1.5 | U266B1 EV | U266B1 sgBcl-2 | NA | NA |
| MCL-1 | Y37 | 160 Gd | Abcam | 2 | U266B1 EV | U266B1 sgMcl-1 | AB_776245 | ab32087 |
| c-MYC | D84C12 | 161 Dy | CST | 2 | TykNU | MCF7 | AB_1903938 | 5605S |
| Iκβα | L35A5 | 164 Dy | CST | 4 | U937 + LPS | U937 | AB_390781 | 4814S |
| BIM | 3C5 | 165 Ho | WEHI (Strasser) | 4 | KMS-12-PE EV | KMS-12-PE sgBIM | NA | NA |
| pRB [S807/811] | J112-906 | 166 Er | BD | 2 | OPM2 + Dex | OPM2 + DMSO | AB_647295 | 558389 |
| pERK 1/2 [T202/Y204] | D13.14.4E | 167 Er | CST | 2 | Ramos + anti-IgM | Ramos | AB_2315112 | 4370S |
| CDK4 | DG93E | 168 Er | CST | 2 | CLL + CD40L | CLL | AB_2631166 | 12790S |
| pH2AX [S139] | JBW301 | 171 Yb | Millipore | 4 | CLL irradiated | CLL | AB_309864 | 05-636 |
| cCaspase3 | C92-605 | 173 Yb | BD | 1 | HELA +TRAIL | HELA - TRAIL | NA | 570524 |
| TP53 | 7F5 | 174 Yb | CST | 2 | Kuromochi | U937 | AB_10695803 | 2527S |
| pS6 [S235/S236] | N7-548 | 175 Lu | BD | 2 | OVCAR4 | U937 | NA | 624092 |
| pCREB [S133] | 87G3 | 176 Yb | CST | 3 | Ramos + anti-IgM | Ramos | AB_2561044 | 9198S |

26

**Supplementary Table 4. Bioinformatic tools used for analysis**

| *Software and Algorithms* | *Identifiers* |
|---|---|
| *Flowjo v10.6.1* | *https://www.flowjo.com* |
| *Cytobank* | *https://www.cytobank.org* |
| *Catalyst* | *https://github.com/HelenaLC/CATALYST* |
| *R v3.4.3* | *https://www.r-project.org/* |
| *CytofRUV* | *https://github.com/mtrussart/CytofRUV* |

**Supplementary Table 5. Murine experimental models**

| Mouse model | Source | Identifier |
|---|---|---|
| *cd23cre* | [43] | MGI:3803652; RRID:IMSR_JAX:028197 |
| $Bak^{-/-}$ | [44] | RRID:MGI:2656013 |
| $cre^{\Delta bax}$ | [45] | RRID:MGI:3589203 |
| *vav-huBcl2* | [46] | RRID:MGI:3842939 |
| $Tnfrsf13c^{-/-}$ | [47] | RRID:MGI:3716689 |
| *Eμ-TCL-1* transgenic | [48] | N/A |

**Supplementary Table 6. Antibodies used for flow cytometric analysis**

| **Antibody** | **Clone** | **Fluorochrome Label** | **Source** | **Dilution** | **RRID** | **Catalogue** |
|---|---|---|---|---|---|---|
| hu CD19 | HIB19 | BV510 | Biolegend | 1:100 | AB_2561668 | 302242 |
| hu CD5 | CLB-T 1/1 | APC | Beckman Coulter | 1:100 | NA | B55386 |
| hu CD4 | OKT4 | BV421 | Biolegend | 1:200 | AB_2562134 | 317434 |
| hu CD19 | HIB19 | BV650 | Biolegend | 1:100 | AB_2562097 | 302238 |
| hu CD5 | UCHT2 | PE-Cy7 | Biolegend | 1:100 | AB_2275812 | 300622 |
| hu BCL-2 | BCL-2/100 | PE-594 | BD | 1:200 | AB_2738307 | 563601 |
| MCL-1 | 19c4-15 | AlexaFluor 647 | WEHI | 1:100 | NA | NA |
| BCL-XL | E18 | PE | Abcam | 1:100 | NA | ab32370 |

| | | | | | | |
|---|---|---|---|---|---|---|
| NOXA | NB600-1159 | AlexaFluor 700 | R&D | 1:200 | NA | NB600-1159AF700 |
| ms CD16/32 | 24G2 | block | WEHI | 1:10 | NA | NA |
| ms CD45.2 | 104 | BUV395 | BD | 1:100 | AB_2738867 | 564616 |
| ms CD43 | S7 | PerCP Cy5.5 | Biolegend | 1:100 | AB_2800667 | 143220 |
| ms CD24 | M1/69 | Pac Blue | Biolegend | 1:100 | AB_572011 | 101820 |
| ms B220 | RA3-6B2 | BV605 | Biolegend | 1:200 | AB_11204069 | 103241 |
| ms CD45.1 | A20.1 | biotin | WEHI | 1:100 | NA | NA |
| Streptavidin | | BV711 | Biolegend | 1:100 | NA | 405241 |
| ms IgM | Nov-41 | BV786 | BD | 1:200 | AB_2741429 | 743328 |
| ms IgD | 11-26c2a | APC-Cy7 | Biolegend | 1:200 | AB_10662544 | 405716 |
| ms Ki-67 | NA | FITC | BD | 1:50 | NA | 665127 |
| ms BCL-2 | 10c4 | PE-Cy7 | eBioscience | 1:100 | AB_2573516 | 25-6992-42 |
| ms CD23 | B3B4 | BUV737 | BD | 1:300 | AB_2873931 | 749668 |
| ms CD21 | 7G8 | BV421 | BD | 1:100 | AB_2737921 | 562966 |
| ms CD25 | PC61 | BV510 | Biolegend | 1:200 | AB_2562270 | 102042 |
| ms CD45.2 | 104 | BV711 | Biolegend | 1:100 | AB_2616859 | 109847 |
| ms CD23 | B3B4 | AlexaFluor 700 | Biolegend | 1:100 | AB_2687125 | 101632 |
| ms CD45.1 | A20 | APC-Cy7 | Biolegend | 1:200 | AB_313505 | 110716 |
| ms CD4 | GK1.5 | PerCP Cy5.5 | Biolegend | 1:200 | AB_893324 | 100434 |
| ms CD184 | 2B11 | BV421 | BD | 1:50 | AB_2737757 | 562738 |
| ms CD45.2 | 104 | BV605 | Biolegend | 1:200 | AB_2563485 | 109841 |
| ms CD8 | 53-6.7 | BV650 | Biolegend | 1:200 | AB_2563056 | 100742 |
| ms CD3 | 17A2 | BV711 | Biolegend | 1:100 | AB_2563945 | 100241 |
| Streptavidin | | BV786 | BD | 1:200 | AB_2869529 | 563858 |
| ms CD44 | IM7 | AlexaFluor 700 | Biolegend | 1:200 | AB_493713 | 103026 |
| ms FoxP3 | FJK-16s | e450 | eBioscience | 1:200 | AB_1518812 | 48-5773-82 |

| | | | | | | |
|---|---|---|---|---|---|---|
| ms CD62L | MEL-14 | APC-Cy7 | Biolegend | 1:200 | AB_830799 | 104428 |
| ms MHC Class II | M5/114. 15.2 | biotin | Biolegend | 1:200 | AB_313731 | 116504? |
| ms Gr1 | RB6-8C5 | biotin | Biolegend | 1:200 | AB_313319 | 107604 |
| ms TER119 | TER-119 | biotin | Biolegend | 1:200 | AB_313369 | 108404 |
| ms CD11b | M1/70 | biotin | Biolegend | 1:200 | AB_313705 | 116204 |
| ms B220 | RA3-6B2 | biotin | Biolegend | 1:200 | AB_312787 | 101204 |
| ms CD11c | N418 | biotin | Biolegend | 1:200 | AB_312989 | 103204 |
| ms TCRgd | GL3 | biotin | Biolegend | 1:200 | AB_313773 | 117304 |

#### Supplementary Figure Legends

**Supplementary Figure 1 (related to Figure 1 and 2). (A)** Structure of the CyTOF runs. All samples were run in 8 batches. Where possible, samples from single patients were grouped together. All runs share reference control samples (PB from a healthy donor and PB from patient CLL25 at 50mg), highlighted in red, for batch effect correction. **(B)** Heatmap summary of median scaled protein expression measure by mass cytometry in each cluster in all cells in peripheral blood (Figure 1C). **(C)** Heatmap summary of median scaled protein expression measure by mass cytometry in each cluster after re-clustering (Figure 2A).

**Supplementary Figure 2 (related to Figure 2). (A)** Violin plots of the mean frequency of all CLL sub-clusters in all 20 patients at screening, 20 mg, 50 mg, 100 mg, 200 mg and 400 mg venetoclax treatment. **(B)** Representative flow cytometry plot showing an increase in the proportions of $CD5^{low}CXCR4^{high}$ cells upon venetoclax dose-escalation in Patient CLL18. **(C)** Violin plots show mean percentages of cells in each gate as illustrated in B.

**Supplementary Figure 3 (related to Figure 3). (A)** Displays of the representative result of 2-D linear discriminant analysis of data from CLL clusters across increasing doses of venetoclax. The vectors indicate the directions of increasing expression for different markers. The major markers associated with the cellular response are apparent, with the dominant effect being higher BCL-2 expression. **(B)** Median level of BCL-2 by CyTOF analysis in all patients undergoing venetoclax dose escalation. (**C**) Lymphocyte counts (cells/L) before and after 200mg of venetoclax in patient CLL15 and CLL21. (**D**) MFI of BCL-2 protein expression by flow cytometry in samples at screening and at post-200mg venetoclax in patient CLL15 and CLL21. (**E**)Venetoclax sensitivity *in vitro* assay in CLL cells from patient CLL15 and CLL21.

**Supplementary Figure 4 (related to Figure 6). (A)** Gating of CD45.2$^{+}$IgM$^{+}$CD21$^{+}$ mature B cells in unmanipulated control mice with the same genotype or cells recovered from venetoclax treated recipient mice. **(B)** Histograms of BCL-2, MCL-1 and BCL-XL protein expression in donor CD45.2$^{+}$ B cells from unmanipulated control mice with the same genotype or from venetoclax treated recipient mice. **(C)** Geometric mean of BCL-2, MCL-1 and BCL-XL protein expression levels in *Bak*$^{-/-}$*Bax*$^{\Delta cd23}$ and *Tnfrsf13c*$^{-/-}$*Bak*$^{-/-}$*Bax*$^{\Delta cd23}$ B cells from control mice or from venetoclax treated recipient mice measured by flow cytometry. Data are representative of three independent experiments with n=3-6 mice per group. For all bar graphs, mean±SEM are shown, and each symbol represents an individual mouse; Student's two-tailed *t* test was used, *, **, *** and **** denotes P<0.05, P<0.01, P<0.005, P<0.001, respectively, n.s. denotes not significant.

**Supplementary Figure 5 (related to Figure 7). (A)** Schematic representation of the liberated ligand hypothesis. **(B)** Serum concentrations of different cytokines analyzed by the Luminex assay in four patients at indicated timepoints.

### Supplementary Figure 1

## A

| Barcode# | Batch 1 | Batch 2 | Batch 3 | Batch 4 | Batch 5 | Batch 6 | Batch 7 | Batch 8 |
|---|---|---|---|---|---|---|---|---|
| 1 | CLL19-Post-20 mg | CLL23 BM-Screening | CLL21-Post-20mg | CLL31-Screening | CLL39 PB-Screening | *CLL13-Screening* | CLL30 BM-Screening | CLL39 BM-Screening |
| 2 | CLL19-Post-20 mg | CLL23 PB-Screening | CLL21-Post-20mg | CLL31-Post-20mg | CLL39-Post-20mg | *CLL13-Screening* | CLL30 PB-Screening | CLL27-Post-400 mg (31 wk) |
| 3 | CLL19-Post-50 mg | CLL23-Post-50 mg | CLL21-Post-50 mg | CLL31-Post-50 mg | CLL39-Post-50 mg | CLL13-Post-20mg | CLL30-Post-20mg | CLL13 BM-Screening |
| 4 | CLL19-Post-100 mg | CLL23-Post-200 mg | CLL21-Post-100 mg | CLL31-Post-200 mg (24 hours) | CLL39-Post-100 mg | CLL13-Post-50 mg | CLL30-Post-50 mg | CLL33-Post-400 mg (18 wk) |
| 5 | CLL19-Post-200 mg | CLL23-Post-400 mg (3 wk) | CLL21-Post-200 mg | CLL31-Post-200 mg | CLL39-Post-200 mg | CLL13*-Post-100 mg | CLL30-Post-100 mg | CLL20 PB-Post-400 mg (51 wk) |
| 6 | CLL19-Post-400 mg (3 wk) | CLL37-Post-20mg | CLL18-Screening | CLL31-Post-400 mg (3 wk) | CLL39-Post-400 mg (3 wk) | CLL13-Post-400 mg (24 hours) | CLL30-Post-200 mg | CLL20 BM-Post-400 mg (51 wk) |
| 7 | CLL19 PB-Post-400 mg (30 wk) | CLL37-Post-50 mg | CLL18-Post-20mg | CLL12-Post-50 mg | CLL33 BM-Screening | CLL13-Post-400 mg (3 wk) | CLL30-Post-400 mg (3 wk) | CLL25-Post-400 mg (39 wk) |
| 8 | CLL25-Screening | CLL37-Post-100 mg | CLL18-Post-50 mg | CLL12-Post-200 mg | CLL33 PB-Screening | CLL15*-Screening | CLL30-Post-400 mg (19 wk) | CLL28-Post-400 mg (30 wk) |
| 9 | CLL25-Post 20mg | CLL37-Post-200 mg | CLL18-Post-100 mg | CLL12-Post-400 mg (3 wk) | CLL33-Post-20mg | CLL15-Screening | CLL27*-Post-20mg | CLL39-Post-400 mg (15 wk) |
| 10 | CLL25-Post 20mg | CLL37-Post-400 mg (3 wk) | CLL18-Post-200 mg | CLL22 BM-Screening | CLL33-Post-50 mg | CLL15-Post-20mg | CLL36-Screening | CLL25 PB-Post-400 mg (45 wk) |
| 11 | **#CLL25-Post-50 mg** | CLL16-Screening | CLL18-Post-400 mg (3 wk) | CLL22-Screening | CLL33-Post-100 mg | CLL15-Post-50 mg | CLL36-Post-20 mg | CLL25 BM-Post-400 mg (45 wk) |
| 12 | CLL25-Post-100 mg | CLL16-Screening | CLL18-Post-400 mg (16 wk) | CLL22-Screening | CLL33-Post-200 mg | CLL15-Post-100 mg | CLL36-Post-50 mg | CLL36-Post-400 mg (22 wk) |
| 13 | CLL25-Post-200 mg | CLL16-Post-20 mg | CLL20-Screening | CLL22-Screening | CLL28-Screening | CLL15-Post-200 mg | CLL36-Post-100 mg | CLL22 BM-Post-400 mg (39 wk) |
| 14 | CLL27 BM-Screening | CLL16-Post-100 mg (24 hrs) | CLL20-Post-20mg | CLL22-Post-20mg | CLL28-Post-20 mg | CLL15 BM-Post-400 mg (39 wk) | CLL36-Post-400 mg (3 wk) | CLL37-Post-400 mg (19 wk) |
| 15 | CLL27 PB-Screening | CLL16-Post-100 mg | CLL20-Post-50 mg | CLL22-Post-50 mg | CLL28-Post-50mg | CLL15 PB-Post-400 mg (39 wk) | CLL39 PB-Screening | CLL22-Post-400 mg (34 wk) |
| 16 | CLL19 BM-Post-400 mg (3 wk) | CLL16-Post-200 mg | CLL20-Post-100 mg | CLL22-Post-100 mg | CLL28-Post-100 mg | CLL35-Screening | CLL15-Screening | CLL23 PB-Post-400 mg (44 wk) |
| 17 | CLL27-Post-50 mg | CLL16-Post 400 mg | CLL20-Post-200 mg | CLL22-Post-200 mg | CLL28-Post-200 mg | CLL35-Post-100 mg | CLL20-Screening | CLL23 BM-Post-400 mg (44 wk) |
| 18 | CLL27-Post-100 mg | CLL16 BM-Post 400 mg (3 wk) | **#CLL25-Post-50 mg** | **#CLL25-Post-50mg** | CLL28-Post-400 mg (2 wk) | CLL35-Post-200 mg | CLL16-Screening | CLL20-Screening |
| 19 | CLL27-Post-200 mg | **#CLL25-Post-50 mg** | CLL28-Screening | CLL18-Post-50 mg | **#CLL25-Post-50 mg** | **#CLL25-Post-50 mg** | **#CLL25-Post-50 mg** | **#CLL25-Post-50 mg** |
| 20 | **#Healthy_981** | **#Healthy_981** | **#Healthy_981** | **#Healthy_981** | **#Healthy_981** | **#Healthy_981** | **#Healthy_981** | **#Healthy_981** |

## B

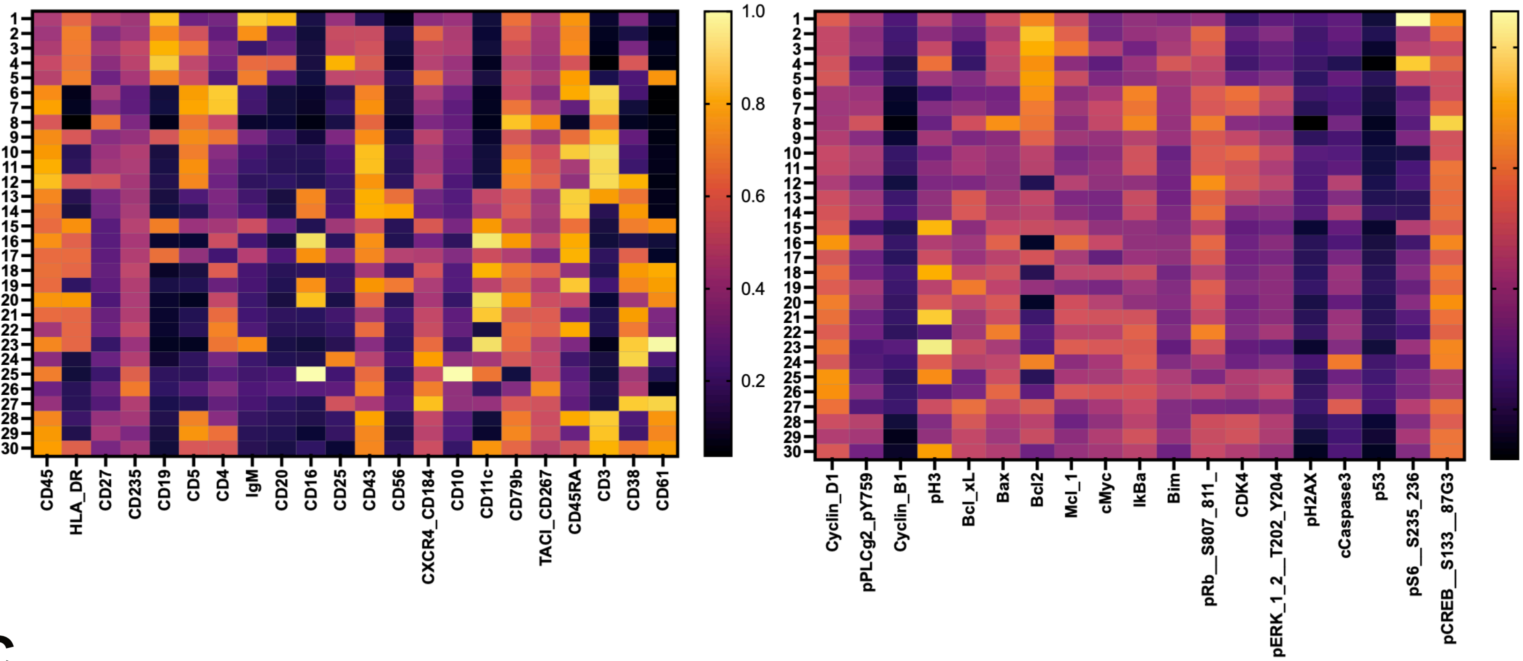

## C

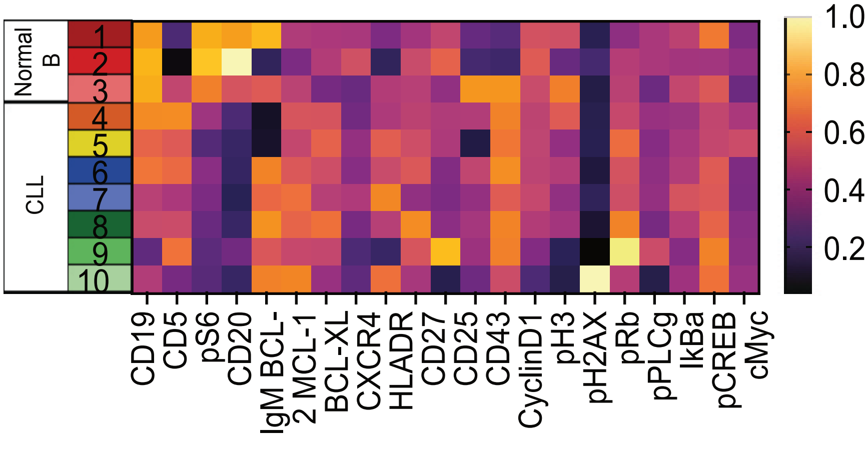

### Supplementary Figure 2

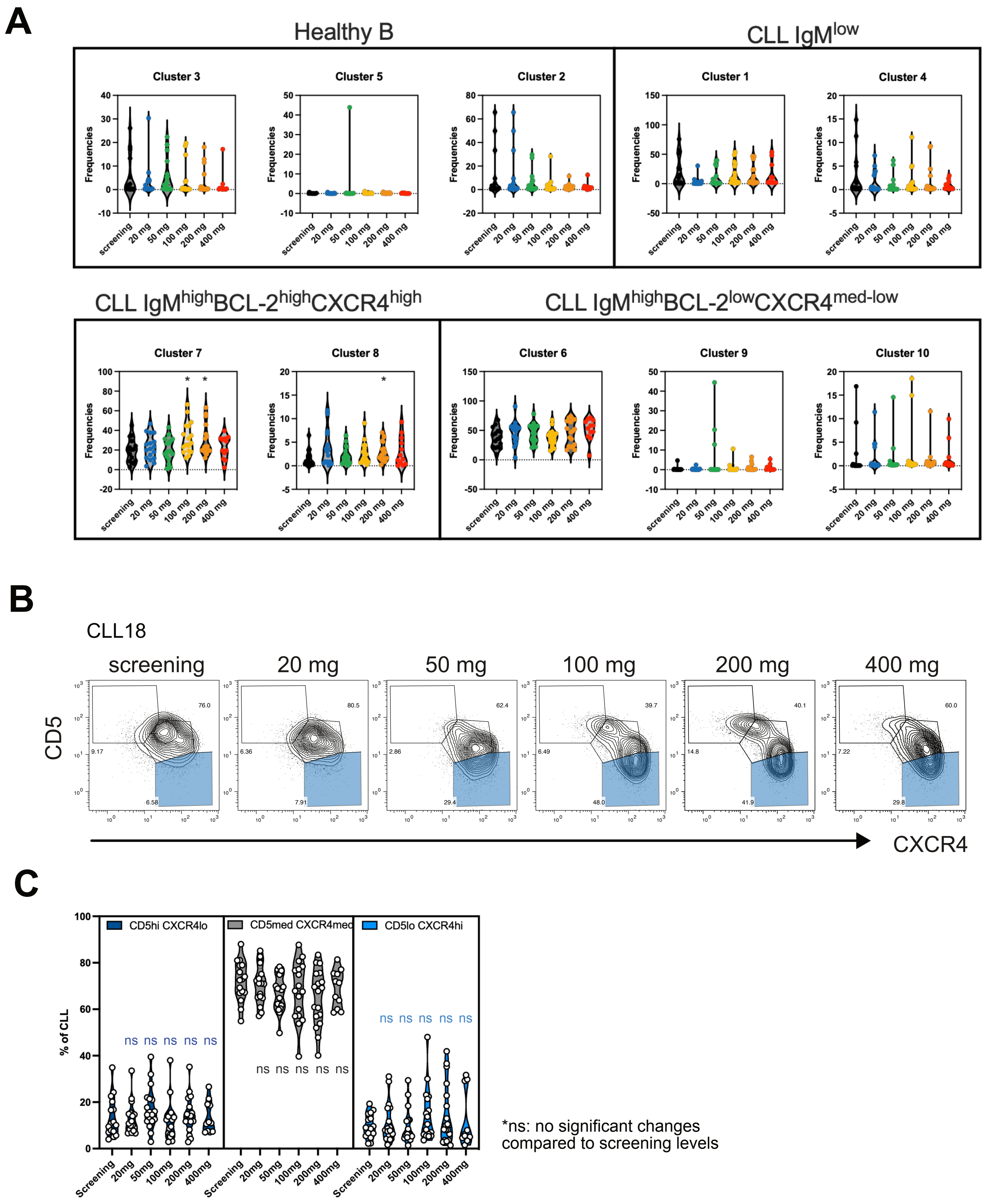

### Supplementary Figure 3

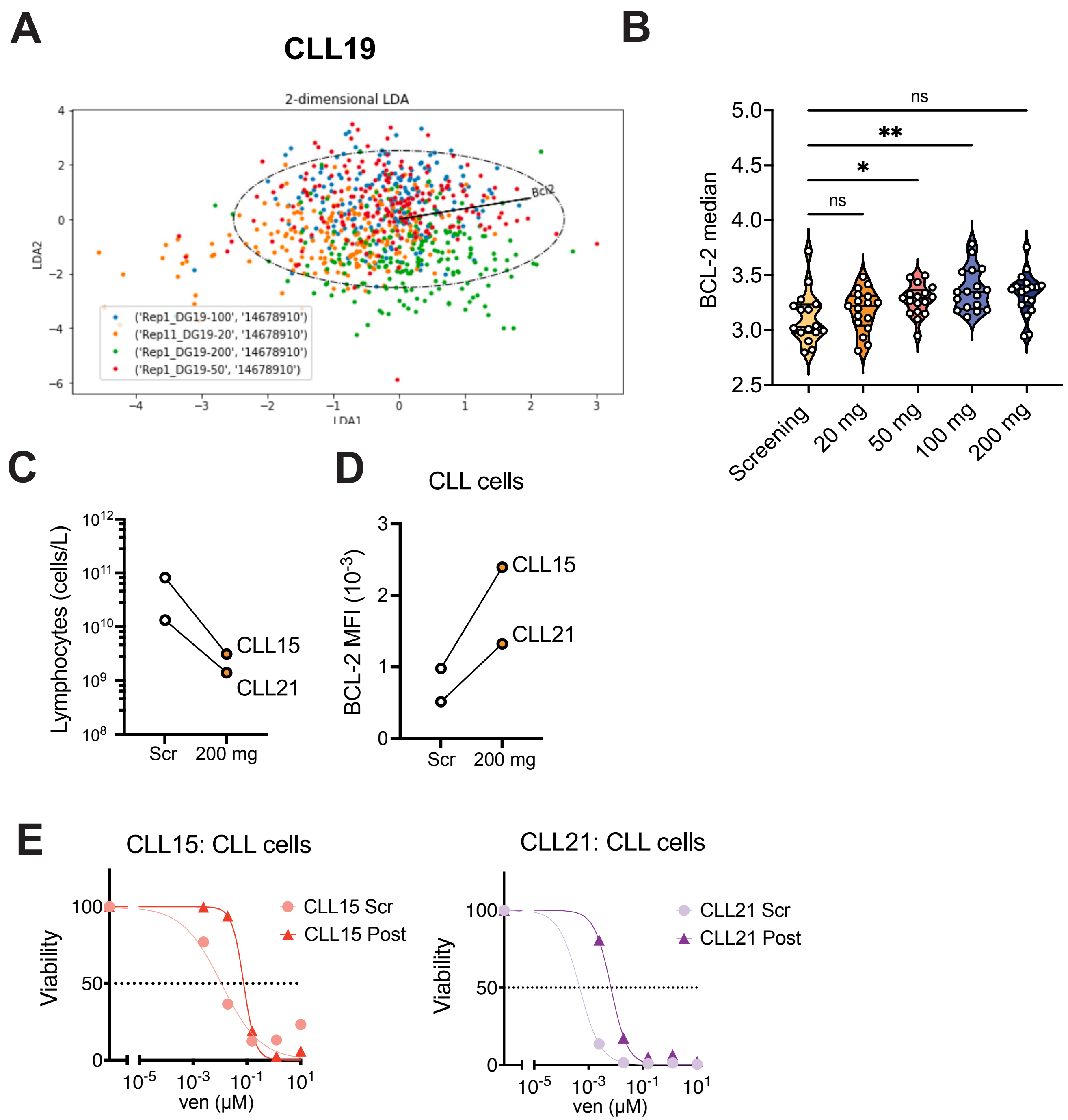

| | DG15 Scr | DG15 Post |
|---|---|---|
| LC50 | 0.01117 | 0.07615 |

| | DG21 Scr | DG21Post |
|---|---|---|
| LC50 | 0.0004699 | 0.006692 |

### Supplementary figure 4

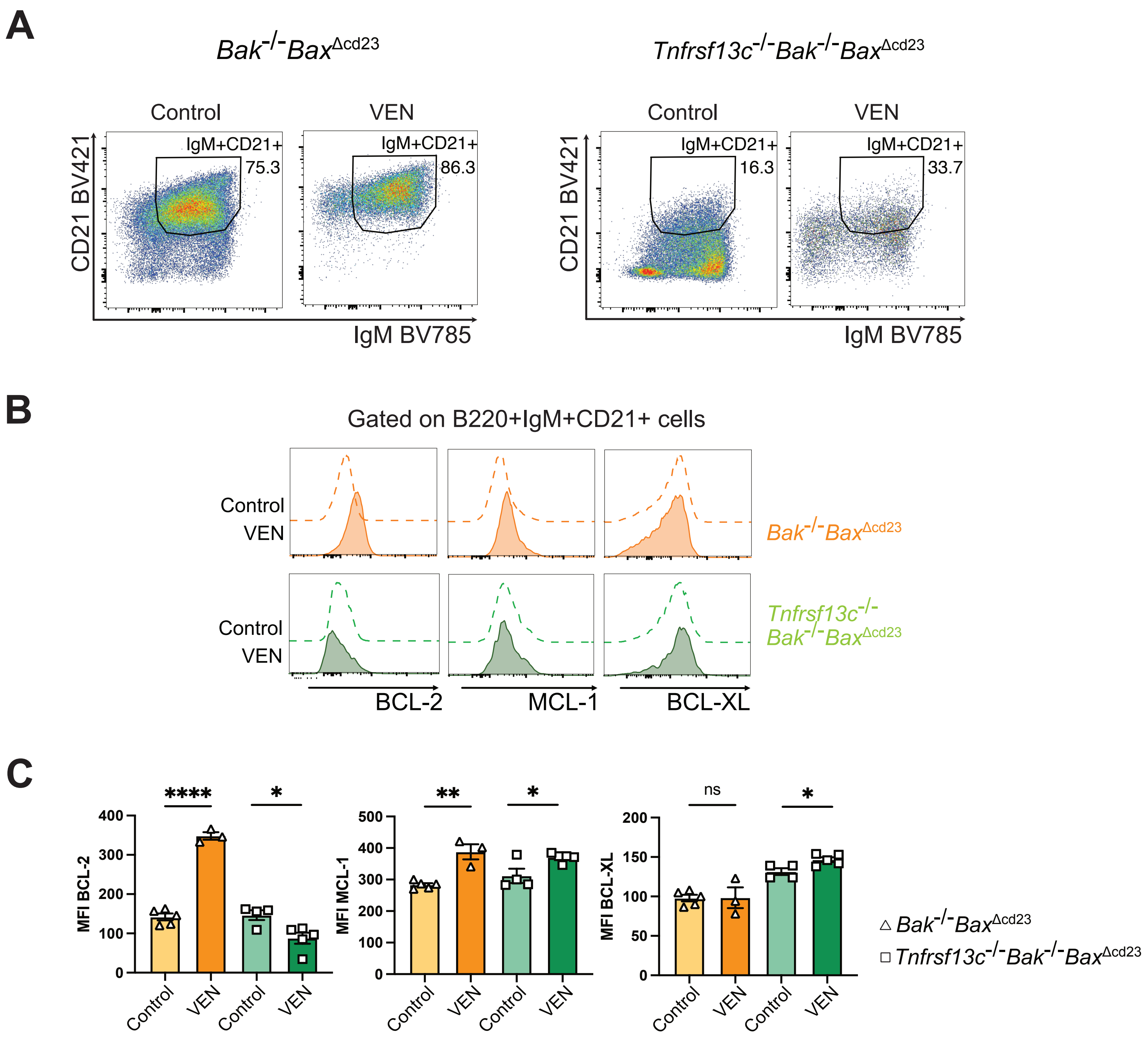

### Supplementary figure 5

A

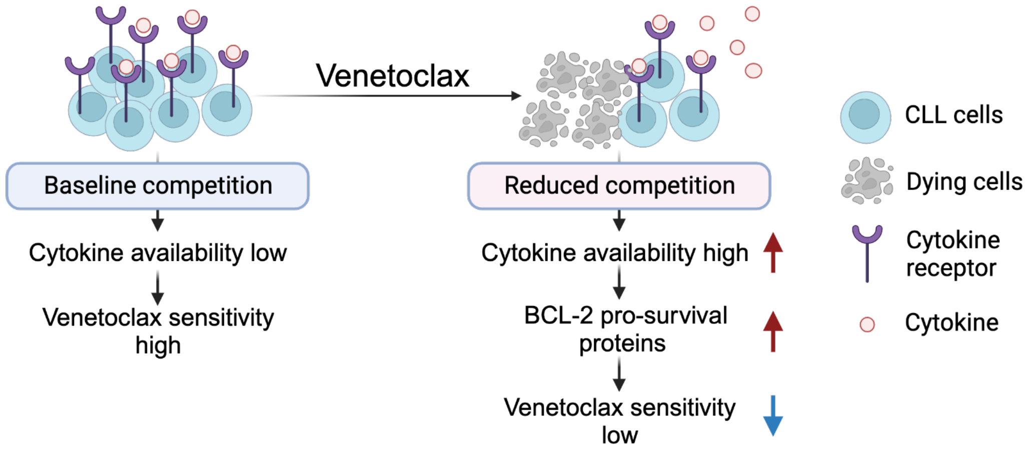

B

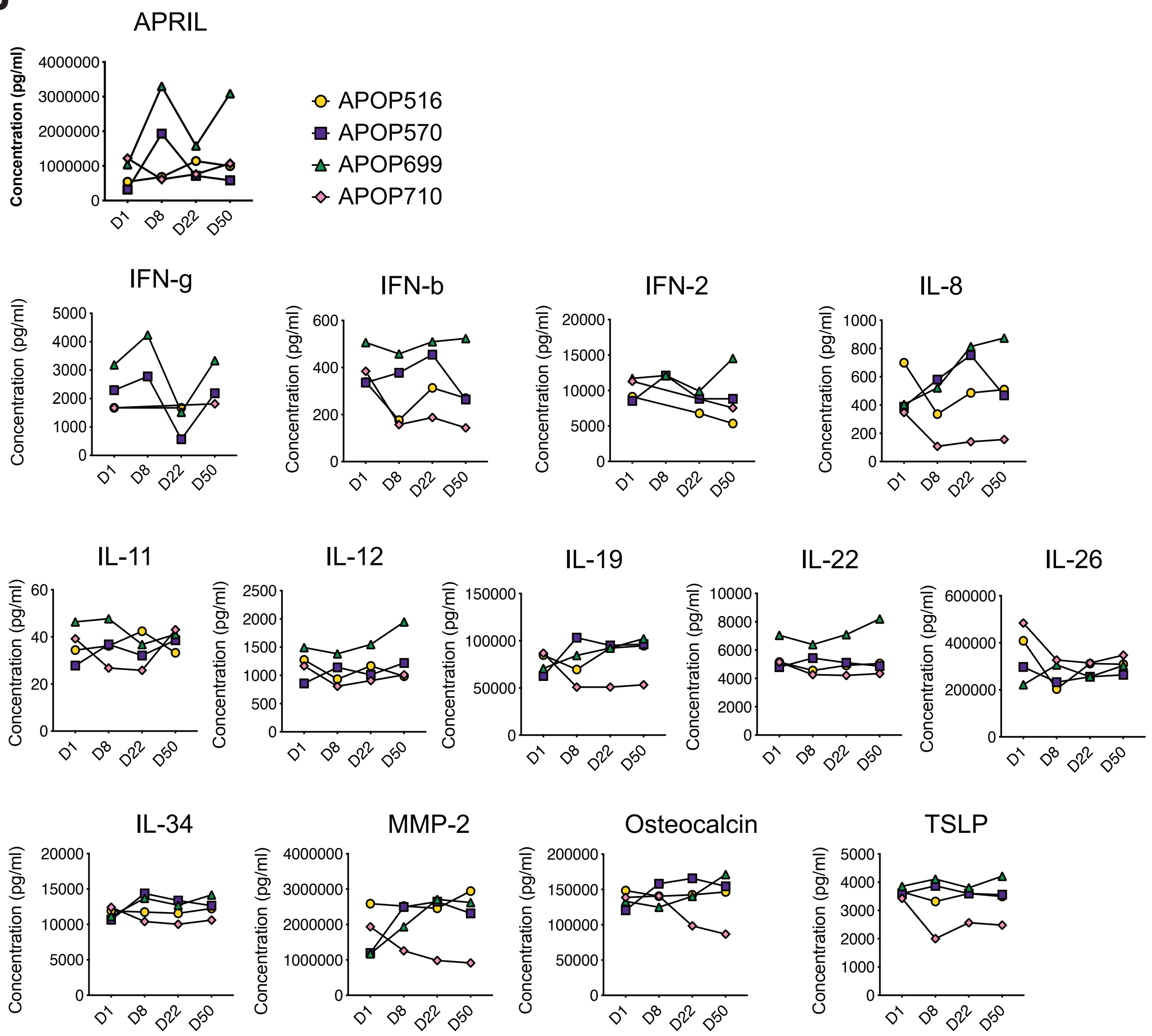
